# Rapid water flow navigates long-distance migration of thermophilic bacteria

**DOI:** 10.1101/2025.02.04.634400

**Authors:** Naoki A. Uemura, Naoya Chiba, Ryota Morikawa, Masatada Tamakoshi, Daisuke Nakane

## Abstract

Bacteria thrive in nearly all environments on Earth, demonstrating remarkable adaptability to physical stimuli, as well as chemicals and light. However, the mechanisms by which bacteria locate and settle in ecological niches optimal for their growth remains poorly understood. Here, we show that *Thermus thermophilus*, a highly thermophilic non-flagellated species of bacteria, exhibits positive rheotaxis, navigating upstream in unidirectional rapid water flow. Mimicking their natural habitat at 70°C with a water current under optical microscopy, cells traveled distances up to 1 mm in 30 min, with infrequent directional changes. This long-distance surface migration is driven by type IV pili, facilitating vertical attachment at the cell pole, and shear-induced tilting of the cell body, resulting in alignment of the leading pole toward the direction of water flow. Direct visualization of T4P filaments and their dynamics revealed that rheotaxis is triggered by weakened attachment at the cell pole, regulated by ATPase activity, which was further validated by a mathematical modeling. Flow experiments on 16 bacterial strains and species in the *Deinococcus-Thermus* phylum revealed that positive rheotaxis is highly conserved among rod-shaped *Thermaceae*, whereas no rheotactic behavior was observed in spherical-shaped *Deinococcus*. Our findings suggest that thermophilic bacteria reach their ecological niches by responding to the physical stimulus of rapid water flow, a ubiquitous feature in hot spring environments. This study highlights unforeseen survival strategies, showcasing an evolutionary adaptation to a surface-associated lifestyle where swimming bacteria would otherwise be swept away.

## Introduction

Microorganisms have evolved diverse adaptive behaviors in response to external stimuli such as light, chemical components, and fluid flow, enabling them to thrive in a wide range of environments (1). The molecular mechanisms underlying bacterial motility encompass both three-dimensional swimming and two-dimensional surface motility (2). In general, however, surface motility in bacteria is slower than flagella-driven swimming. This raises the question: is surface motility inherently insufficient for long-distance migration?

Rheotaxis, directed movement against water flow, is a common response in both eukaryotes and bacteria (3). For instance, human sperm and unicellular eukaryotes such as *Tetrahymen*a swim upstream along solid surfaces in flowing environments (4, 5). Similarly, *Pseudomonas aeruginosa*, a pathogenic bacterium, harbors two distinct motility systems: flagella and type IV pili (T4P). In surface-associated lifestyles, T4P-dependent motility is predominant, albeit slower than flagellar motility, allowing bacteria to move against water flow (6). Certain small bacteria, such as *Mycoplasma*, exclusively rely on gliding motility, specialized for movement on solid surfaces, allowing them to move against rapid flow (7, 8). The fundamental basis of these rheotactic responses involves the orientation of the cell body parallel to the flow direction due to shear forces at the liquid-solid interface. However, as the size of the microorganisms decreases, the shear stress required for the orientation increases. In a fast-flowing environment where shear stress exceeds 10 mPa, small life forms such as bacteria are expected to encounter limitations in free swimming (9), and may also struggle to detect chemical gradients (10). *Thermus thermophilus* is a rod-shaped, non-flagellated thermophilic species of bacteria first isolated from a hot spring in 1968 (11), where water flow is a ubiquitous feature (12). Due to its remarkable adaptability to high temperatures, this species serves as a model organism in the research of protein structure, biotechnological applications, and extremophile biology (13–16). *T. thermophilus* harbors T4P machinery for surface motility as well as other processes, including surface attachment, biofilm formation, DNA uptake, and phage infection, (17). In rod-shaped bacteria, T4P machinery are typically localized at the cell poles, where the activity is dynamically controlled to enable cellular movement through repeated cycles of pilus extension and retraction, driven by ATPase motors (17). The protein components of T4P are conserved in the *Thermaceae* family and are closely related to those found in the T4P components of *Deinococcus* (18), a genus renowned for its extraordinary resistance to radiation (19). This conservation implies that T4P plays a critical role in the survival strategies of the *Deinococcus-Thermus* phylum. However, *T*. *thermophilus* lacks a conventional chemotaxis gene set (20), leaving the functional significance of its T4P-dependent motility machinery largely unexplored.

Here, we designed an experimental system that mimics the natural water flow conditions encountered by *T. thermophilus* in its natural environment, allowing us to visualize its directional upstream movement over surfaces. Single-cell imaging of T4P filaments and dynamics reveals that water flow recognition is likely governed by the coordinated activity of dual T4P motors. Furthermore, water flow experiments involving 15 strains and species from the *Deinococcus-Thermus* phylum demonstrate that positive rheotaxis is characteristic in *Thermus* and related thermophilic bacteria. These findings suggest that *T. thermophilus* and its relatives have adapted to fast-flowing environments, where flagellated bacteria are unable to achieve effective translocation.

## RESULTS

### Upstream movement of *T. thermophilus* induced by poor nutrients

The motility of *T. thermophilus* has previously been demonstrated as colony migration on agar plates (21). To observe its motility under an optimal growth temperature at 70℃, we setup a stage heater integrated with an optical microscope to visualize single-cell movement. On the glass surface, *T. thermophilus* HB8 cells showed random motility with a net displacement close to zero for 1 min (Fig 1A *Leftmost* and Movie S1). The apparent diffusion coefficient of the cells was 1.7 µm^2^/s, consistent with the T4P-dependent motility observed in other bacteria (22, 23).

**Fig 1.**
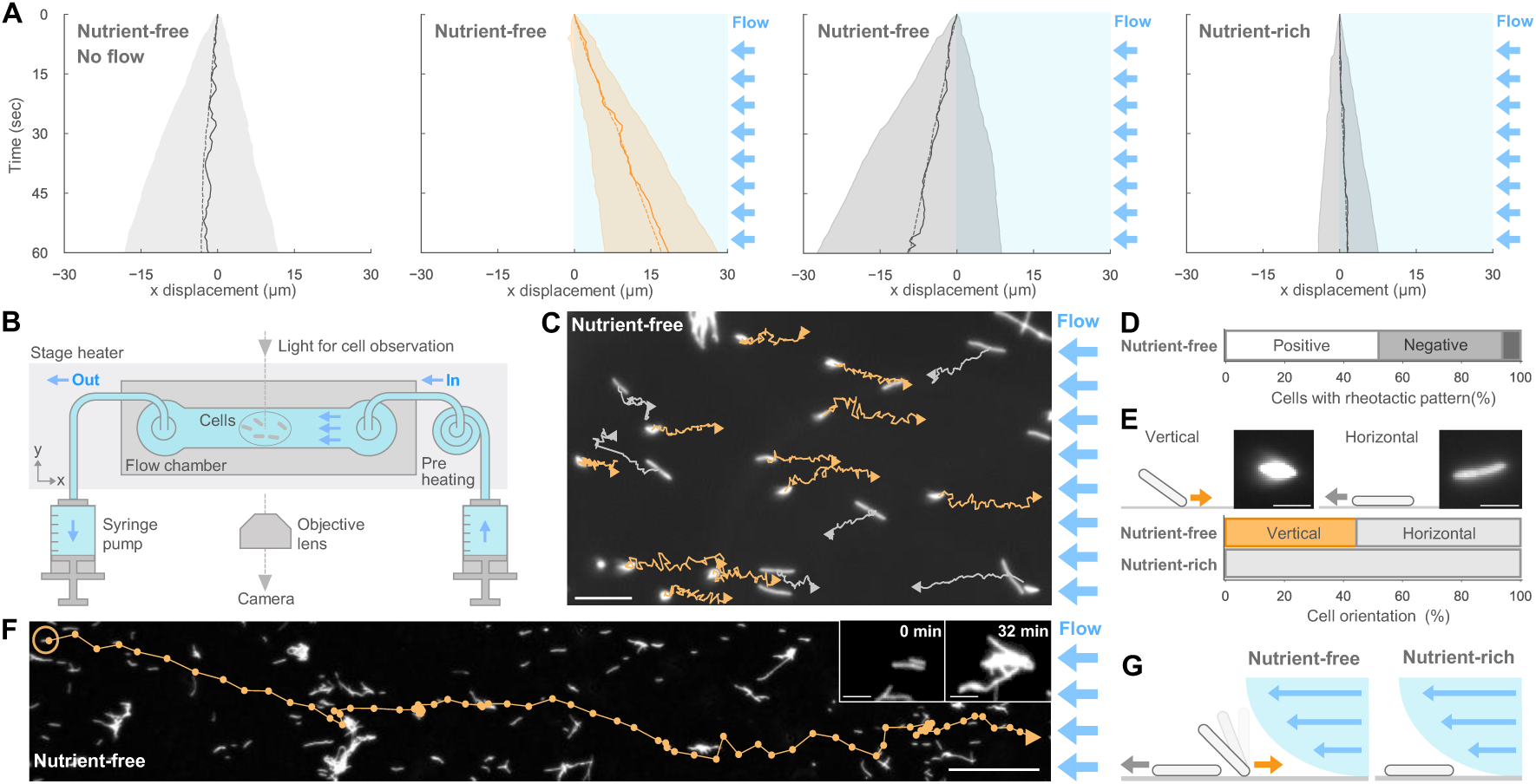
Rheotaxis of *T. thermophilus*. (A) Time course of cell displacements in response to water flow. Dash and solid lines show the average and the median displacement, respectively. The shaded area indicates the standard deviation (SD) of biological replicates (n = 50 cells). (B) Schematic diagram of the experimental setup. Water flow was applied from the right side of the fluid chamber using a syringe pump. Cell behaviors were observed at 70°C under an optical microscope. (C) Field view of cell behavior under a water flow of 40 µm/s in nutrient-free conditions. Cell trajectories for 1 min are overlayed on a dark-field microscopy image captured at time zero. Yellow and grey lines indicate trajectories of vertical and horizontal cells, respectively. Scale bar, 20 µm. (D) Proportion of cell behaviors in water flow (n = 81 cells). White: positive rheotaxis (displacement ≥ 1 µm/min with trajectory angles ≤ 60° relative to upstream direction). Dark grey: negative rheotaxis (displacements for ≥ 1 µm/min with trajectory angles ≥ 120°). Black: random motility (other trajectories). (E) Cell orientation under water flow. *Upper*: dark-field images of vertical and horizontal cells. Scale bar, 5 µm. *Lower*: Proportion of cell orientation in nutrient-free (n = 45 cells) and nutrient-rich condition (n = 65 cells). Vertical cell: Surface attachment via one pole for ≥ 10 s. Horizontal cell: Longitudinal cell axis lying near the surface. (F) Single cell trajectory during positive rheotaxis. The trajectory for 78 min at 1-min intervals is overlaid on the image captured at time zero. The start position is marked by a yellow circle. Scale bar, 100 µm. Inset: cell aggregation initiated by horizontal cells in 32 min. Scale bar, 10 µm. (G) Schematic of cell behavior under water flow in nutrient-free and nutrient-rich conditions.

To apply water flow, we constructed a fluid chamber connected to a syringe pump, placed on a stage heater to maintain a constant water flow temperature at 70°C (Fig 1B and Fig S1A). The syringe pump provides precise control of flow rates, and the resultant flow velocity was calibrated using the linear relationship between the flow rate and the initial velocity of cells detached from the chamber surface (Fig S1B). This setup was used in subsequent experiments.

When exposed to water flow at 40 µm/s, corresponding to a shear stress 0.4 Pa, approximately 50% of *T. thermophilus* HB8 cells showed positive rheotaxis, moving upstream on the glass surface, while about 40% showed negative rheotaxis, moving downstream (Fig 1CD and Movie S2). Positive rheotactic behavior was predominant in vertically aligned cells, with their longer axis perpendicular to the glass surface. These vertical cells positioned one pole against the direction of flow, while the other pole swung freely without surface attachment (Fig 1E *Upper left*). These cells exhibited unidirectional upstream movement with a net displacement of 17.1 ± 11.0 µm/min (Fig 1A *Second from Left*). In contrast, negative rheotaxis was observed in horizontally aligned cells, with their longer axis parallel to the surface (Fig 1E *Upper right*). These horizontal cells moved rectilinearly in the same direction of water flow, with a net displacement at −9.2 ± 18.4 µm/min (Fig 1A *Third from Left*). During positive rheotaxis, the cells exhibited a remarkably low reversal rate, with one reversal every 100 minutes. Additionally, the cells traveled upstream more than 1 mm distance without detaching from the surface, maintaining motility for over an hour under water flow at 40 µm/s (Fig 1F and Movie S3). Cell aggregation was sometimes observed within 30 minutes, but was initiated by horizontal cells (Fig 1F).

Notably, positive rheotaxis was predominantly observed under nutrient-free buffer. When we applied the water flow containing growth medium, vertical cells quickly reoriented to a horizontal position within one minute (Fig 1E *Lower*), and rheotaxis was completely abolished, as evidenced by a net displacement approaching zero (Fig 1A *Rightmost* and Movie S4). These results suggest that *T. thermophilus* HB8 actively regulates its vertical orientation on the solid surfaces to optimize its response to water flow under oligotrophic conditions (Fig 1G).

### Rheotaxis of *T. thermophilus* depends on T4P machinery

T4P filaments are known to facilitate vertical orientations of the cell body in other bacterial species (24, 25). To test whether the polar attachment in *T. thermophilus* HB8 is mediated by T4P machinery, we examined a Δ*pilA* mutant, which lacks the TTHA1221 gene that encodes the major pilin. The Δ*pilA* mutant displayed no T4P filaments and maintained horizontal orientations on the glass surface (Fig 2AB, and Fig S2), indicating that T4P filaments are essential for transitioning to polar attachment. To further dissect the mechanism underlying this transition, we focused on the two putative ATPases of the T4P machinery: PilT1 and PilT2. Generally, PilT1 acts as the primary motor for pilus retraction, while PilT2 is considered a paralog of PilT1 (21, 26). In *T. thermophilus* HB8, a single-deletion mutants Δ*pilT1*, as well as the double-deletion mutant Δ*pilT1*Δ*pilT2*, completely lost motility (Fig 2BC and Fig S3). Both of these mutants, however, exhibited both horizontal and vertical orientations under flow (Fig 2AB). Interestingly, while Δ*pilT2* mutant lost most of its rheotactic capability (Fig 2C), a few vertical cells exhibited weak rheotaxis several minutes after the onset of water flow, with a net displacement 10 times lower than that of the WT (Fig 2D and Movie S5). These data demonstrate that PilT1 functions as a primary motor for T4P-driven cell propulsion, consistent with previous reports (21, 26), while PilT2 appears to play a secondary role that contributes to rheotaxis and enhances sensitivity to water flow.

**Fig 2.**
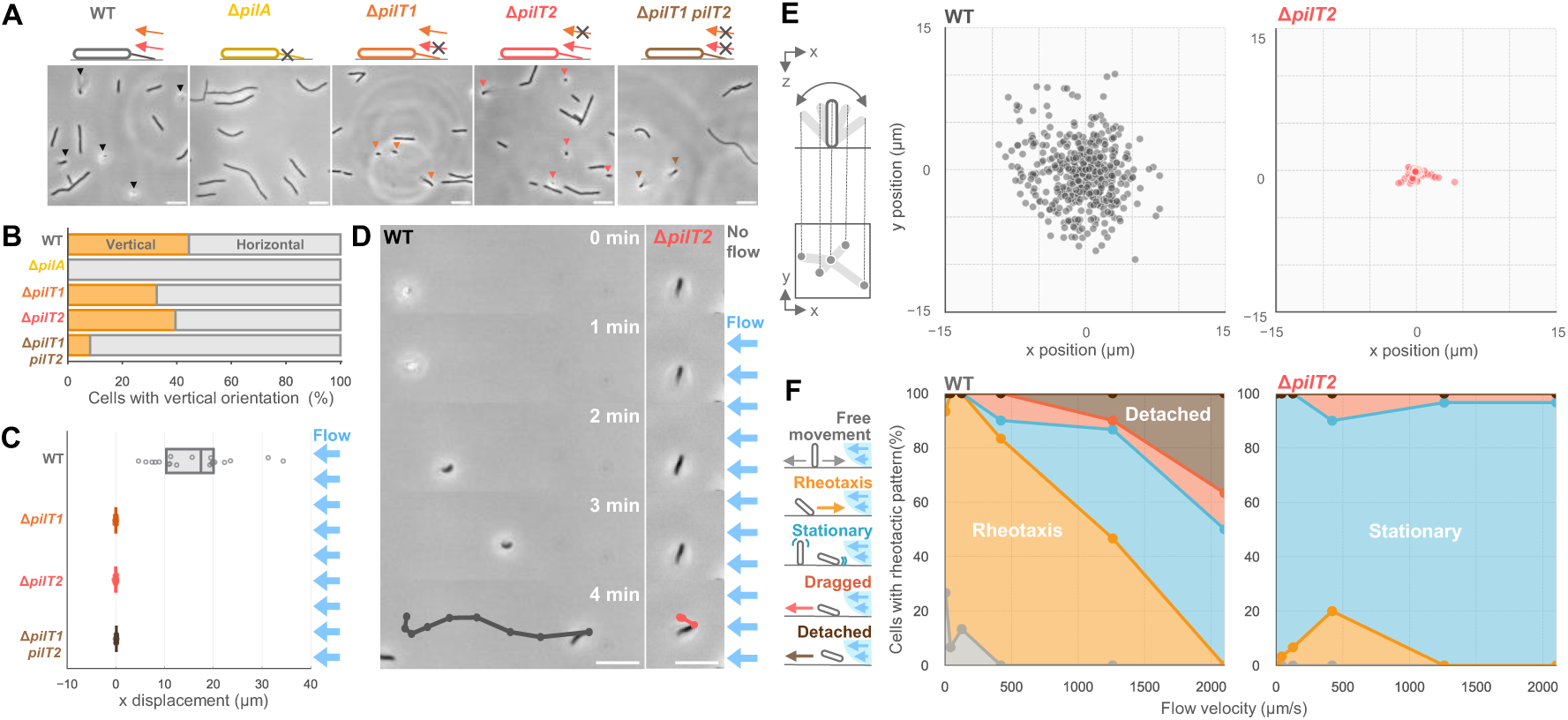
Rheotaxis depends on T4P machinery. (A) Phase-contrast images of *T. thermophilus* WT and T4P mutants. Triangles indicate vertical cells. Scale bar, 10 µm. (B) Proportion of cell orientation in T4P mutants. n = 45 (WT), 60 (Δ*pilA*), 92 (Δ*pilT1*), 75 (Δ*pilT2*), 85 (Δ*pilT1pilT2*) cells. Classifications follow Fig 1E. (C) Rheotaxis velocity of vertical cells in T4P mutants at a water flow of 40 µm/s. Positive values indicate upstream movement. Circles represent biological replicates, and boxplots show the median with 25%/75% quantiles. n = 20 (WT, Δ*pilT1,* and Δ*pilT2*) and 15 (Δ*pilT1pilT2*) cells. (D) Time-lapse images of WT and Δ*pilT2* cells under water flow. Cell trajectories for 4 min at 30-s intervals are overlaid on the last image. Water flow was applied from the right side starting at 1 min. Scale bar, 10 µm. (E) Flapping motion of vertical cells in WT and Δ*pilT2* without water flow. *Left*: Schematic of flapping motion analysis, where the unattached pole position is determined relative to the attached pole. *Right*: Distribution of the unattached pole position for 30 s (n = 450 from 15 cells at 1-s intervals). (F) Proportion of responses to water flow at varying velocities in vertical cells of WT and Δ*pilT2*. Responses are classified into five categories in the left schematics. Rheotaxis: upstream displacement ≥ 1 µm/min within trajectory angles ≤ 60°. Dragged: downstream displacement ≥ 1 µm/min within ≥ 120°. Detached: downstream displacement ≥ 13 µm/min within ≥ 120°. Stationary: displacement ≤ 1 µm/min along flow direction and ≤ 5 µm/min perpendicular. Detached: 3-min displacement for ≥ 3 µm along, and ≤ 10 µm perpendicular. Free movement: other trajectories. n = 30 cells at each flow velocity.

### Sensing flow direction through cell body flapping

Previous studies have demonstrated that polar attachment is required for T4P-dependent rheotaxis in *Pseudomonas* (6). We speculated that vertical orientation of *T*. *thermophilus* cells is critical for sensing water flow near the surface. Prior to the introduction of water flow, vertical cells exhibited a flapping motion around a small area without stable attachment (Fig 2E *Left* and Movie S6). Upon the application of water flow, these cells experienced shear forces that aligned their longer axis parallel to the direction of flow. Subsequently, the cells oriented their leading pole against the flow direction and moved upstream (Fig 2D *Left*). On the other hand, Δ*pilT2* mutant cells maintained their upright orientation but remained immobilized (Fig 2E *Right* and Movie S6), irrespective of whether water flow was present or absent (Fig 2D *Right*). To quantify the difference in adhesion, we calculated the apparent spring constant of vertical cells, assuming that the flapping motion was driven by polar attachment (Fig S4). Notably, the apparent spring constant of the rheotaxis-defective Δ*pilT2* mutant was measured as 3.6×10^−2^ pN/nm, approximately 500 times higher than that of WT (5.9×10^−5^ pN/nm). These results suggest that *T*. *thermophilus* weakens its surface attachment during vertical orientation by PilT2, enabling the cell to effectively sense flow direction.

To validate this hypothesis, we examined the behavior of vertical cells under higher flow velocities (Fig 2F). WT cells showed a reduction in positive rheotaxis as flow velocity increased, ultimately reaching zero at a velocity of 2,000 µm/s. At this threshold, WT cells also began detaching from the glass surface. In contrast, the Δ*pilT2* mutant cells showed minimal positive rheotaxis but remained attached to the surface, even at higher flow velocities. We quantified the maximum forces for attachment based on the drag coefficient of the cell body and the corresponding flow velocity (Fig S5). The maximum forces were estimated to be 132 ± 17 pN for WT cells and 337 ± 17 pN for Δ*pilT2* mutants, indicating that deletion of *pilT2* significantly enhances T4P-dependent surface attachment. These observations suggest that PiT2 activity appears to facilitate the weakening of surface attachment, implying that a balance between surface attachment and motility is essential to adapt to flowing environments.

### Visualization of T4P filaments

We sought to determine whether the differences in surface attachment between WT and the Δ*pilT2* mutant depend on the localization and dynamics of T4P filaments. Using immunofluorescence microscopy of PilA, T4P filaments were directly visualized in vertical cells (Fig 3A *Left*). WT cells showed asymmetrically distributed T4P filaments, predominantly extending from the side opposite to the water flow. The average filament length was 2.8 ± 3.0 µm, and the visible number was 7.6 ± 2.8 per cell (Fig 3B *Left*, and Fig S6). In contrast, Δ*pilT2* mutant showed radially distributed T4P filaments (Fig 3A *Right*), with an increased average length of 4.5 ± 4.4 µm and a significantly higher visible number of 13.4 ± 5.4 per cell (Fig 3B *Right*, and Fig S6). These observations demonstrate that rheotaxis is driven by the asymmetric distribution of T4P filaments, which is disrupted in the absence of PilT2. To further understand the difference between vertical and horizontal cells, we visualized T4P filaments in horizontal cells in the WT. T4P filaments were distributed at both poles, but T4P filaments were observed with high density at the leading pole (Fig S7ABC). This polar localization pattern was similarly observed in Δ*pilT2* mutant cells, consistent with the observation in vertical cells (Fig 3B and Fig S6B). In addition, non-moving Δ*pilT2* cells showed a significantly higher number of T4P filaments compared to WT (Fig S7DE). This suggests that the reduced rheotactic sensitivity in Δ*pilT2* mutants may result from altered rates of T4P extension and retraction.

**Fig 3.**
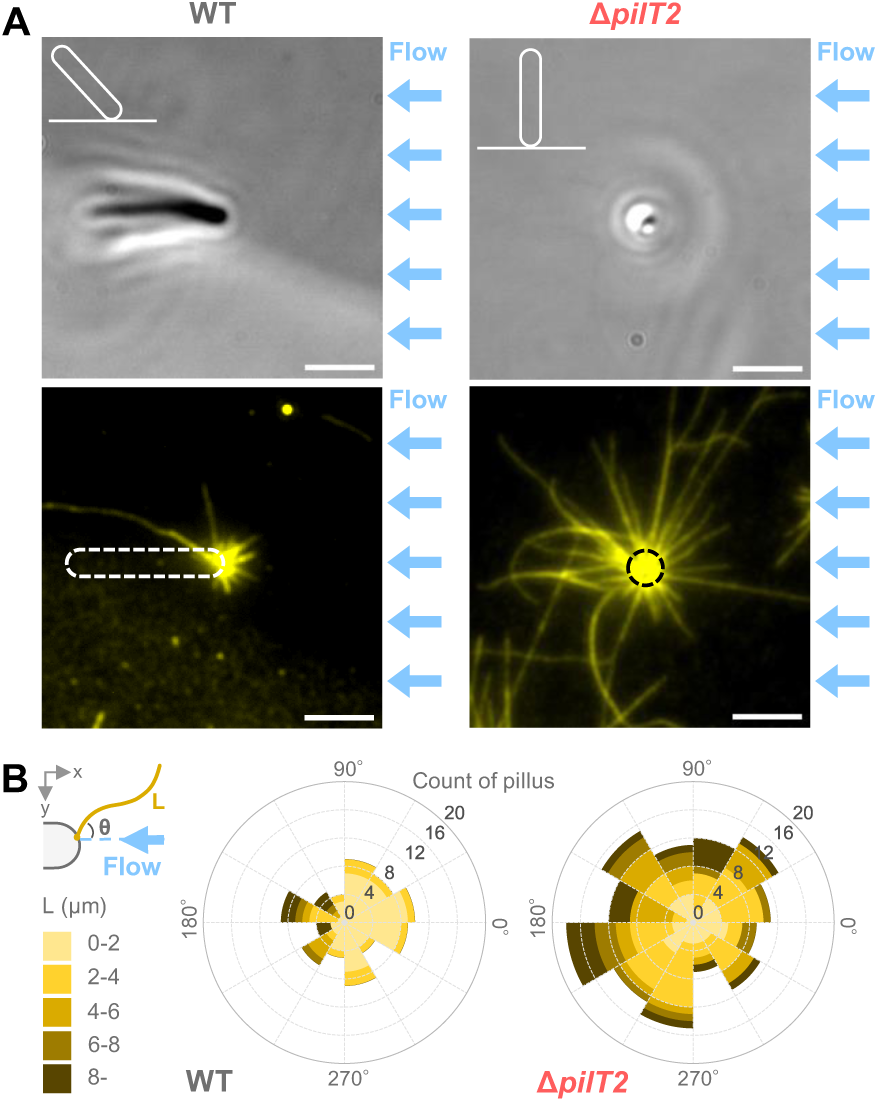
Visualization of T4P filaments by immunofluorescence microscopy. (A) Phase-contrast (top) and PilA immunofluorescence (bottom) images of vertical cells in WT and Δ*pilT2*. Cells were chemically fixed under water flow applied from the right in nutrient-free conditions. White and black dashed lines outline the cell body. Schematic of cell orientations are shown in the upper left of each image. Scale bar, 3 µm. (B) Length and angular distribution of T4P filaments in vertical cells. Left: Schematic image defining filament angle θ relative to the flow direction and filament length L. Right: rose plot displays the distribution of T4P filaments, and lengths in WT and Δ*pilT2* (n = 11 cells for each strain).

### Visualization of T4P dynamics

We visualized T4P dynamics to track the movement of 200 nm silica beads attached to T4P filaments (Fig 4A), as described in previous studies in other bacterial species (23, 27). Beads near the cell pole showed directional movements at constant velocities (Fig 4BC and Movie S7). In WT cells, the average velocity of beads toward the leading pole was measured as −3.04 ± 0.81 µm/s, while the velocity away from the leading pole was 1.01 ± 0.62 µm/s (Fig 4D *Top*). Retraction of beads was also observed at the lagging pole, with comparable velocity almost at both poles, indicating that the number of T4P filaments determines the polarity of cell migration (Fig S8 and Movie S8). In Δ*pilT2* mutant cells, the average velocity of beads movements toward and away from the leading pole significantly reduced to −1.27 ± 0.41 µm/s and 0.65 ± 0.57 µm/s, respectively (Fig 4D *Bottom* and Movie S9). Additionally, the frequency of directional bead movement was lower in Δ*pilT2* mutants compared to WT cells (Fig 4E). Bead dynamics such as the fractions of extension and retraction events, however, remained similar between WT and Δ*pilT2* (Fig S9).

**Fig 4.**
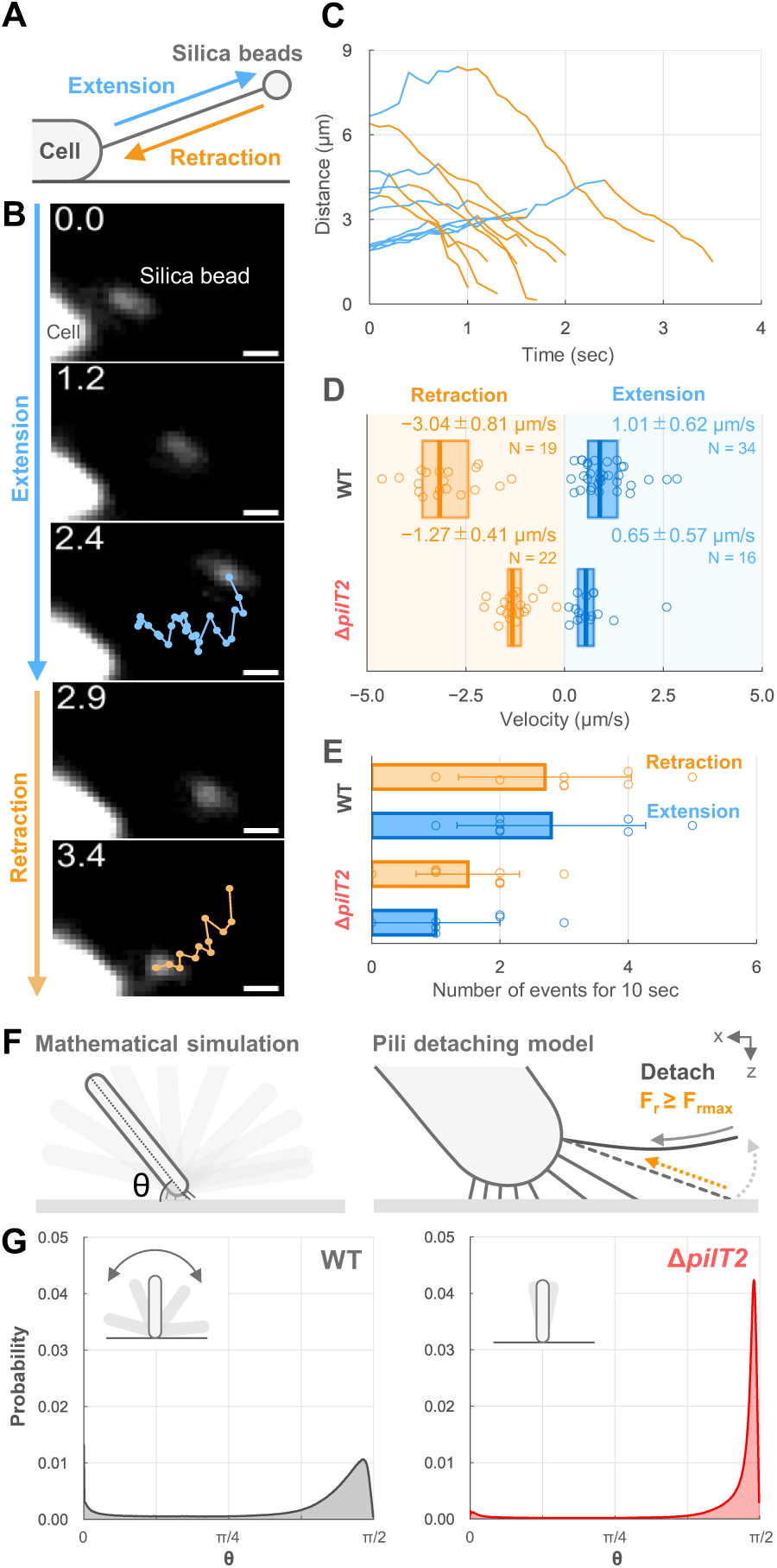
Visualization of T4P dynamics via nanobeads. (A) Schematic of the bead assay. T4P dynamics were visualized using fluorescent silica beads attached to T4P filaments. Bead movements were observed near the leading pole of horizontal cells in nutrient-free conditions. (B) Bead trajectory, captured under dark-field microscopy, showing movement away from and toward the cell pole for 3.4 s. Scale bar, 1 µm. (C) Time course of bead displacement relative to the cell pole in WT. Blue and orange lines represent bead movements away from and toward the cell pole (n = 12 beads from 12 cells). (D) Velocity of bead movement in WT and Δ*pilT2*. Positive values show movement away from the cell pole. Directional bead movements for more than 0.5 s were used for data analyses. Circles indicate biological replicates, and boxplots display the median and 25%/75% quantile (n = 8 cells for each strain). (E) Frequency of directional bead movements in WT and Δ*pilT2.* Frequency was measured for 10 s after the first event. Circles indicate biological replicates, and bars represent average and SD (n = 10 cells for each strain). (F) Schematic of mathematical simulation. Left: angle between the cell body and the surface over time is calculated. See more details in *SI Appendix,* Supplemental Text, and Tables S1 and S2. Right: the model of simulation assumes pili detach when the applied force exceeds the retraction force limit F_rmax_. (G) Angle distribution from the mathematical simulation in WT and Δ*pilT2.* Data represent n = 5 × 10^6^ from 500 cells over 1,000 s at 0.1-s intervals. Insets show schematic of cell behaviors.

### Dual motor controls vertical orientation for rheotaxis

We hypothesized that the coexistence of dual PilT motors enhances the flapping motion of vertical cells, contributing to rheotaxis. To test this hypothesis, we developed a pili detachment model based on a tug-of-war mechanism (22), where multiple T4P filaments coordinate their activity through force-dependent detachment (Fig 4F *Right,* see more details in Supplemental Text). Assuming that T4P filaments extend from one pole, we simulated the behavior of vertical cells using parameters of T4P distribution and dynamics (Fig 4F *Left,* Tables S1 and S2). Simulations based on Δ*pilT2* parameters predicted that cells would maintain an upright orientation, strongly anchored at a single pole, with minimal fluctuation in cell angle (Fig 4G *Right*). In contrast, simulations with WT parameters predicted that vertical cells would occasionally tilt with substantial fluctuations in cell angle (Fig 4G *Left*). Our results suggest that the flapping motion is driven by a tug-of-war competition, which is accelerated by the diverse retraction activity in the presence of dual motors.

### Responses to water flow in related species

We considered the prevalence of rheotactic behavior in closely related species. Based on the criteria that bacteria harbor putative genes for T4P machinery and two or more *pilT* homologs in their genomes, we selected six strains of *T. thermophilus* and ten additional species within the *Deinococcus-Thermus* phylum (Fig 5A and Table S3). All bacteria were cultured and observed under their optimal growth temperature using optical microscopy. Surface motility was seen in 14 bacterial strains and species, including *Thermus aquaticus, Meiothermus ruber,* and *Deinococcus radiodurans* (Movie S10). Single-cell trajectories revealed that surface motility was diffusive in the absence of the water flow, consistent with our observation of *T. thermophilus* HB8 (Fig 5B and Fig S10). *Deinococcus radiophilus*, however, was barely motile and excluded from subsequent experiments.

**Fig 5.**
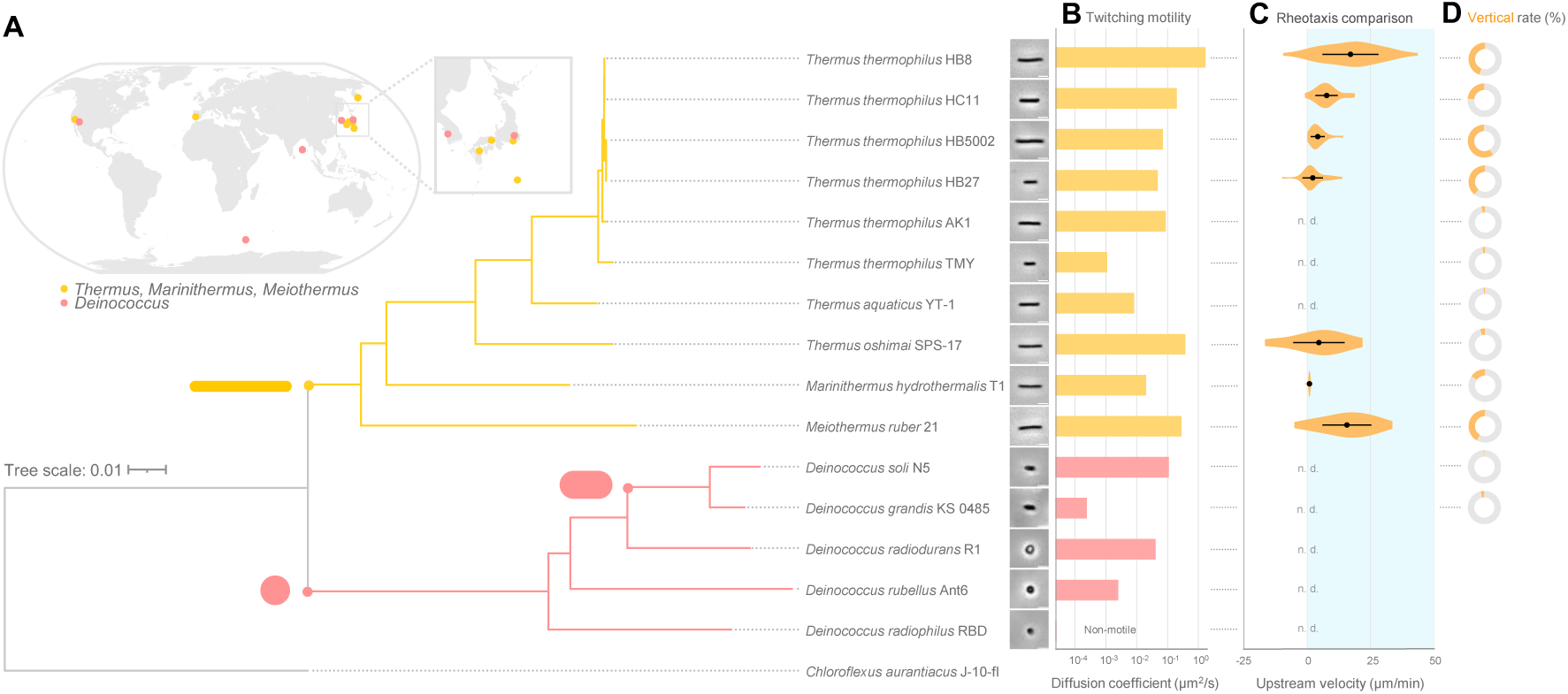
Twitching motility and rheotaxis in *Deinococcus-Thermus*. (A) Phylogenetic tree and isolation locations of *Deinococcus-Thermus*. *Upper left*: world map showing the isolation site of each strain. *Right*: the phylogenetic tree is based on 16S ribosomal RNA sequences with *Chloroflexus aurantiacus* as an outgroup. Yellow and red branches correspond to *Thermaceae* and *Deinococcus*, respectively. Phase-contrast images of each strain are shown. Scale bar, 3 µm. (B) Twitching motility assay. Diffusion coefficient of the cell movements over glass surfaces in the absence of water flow (see more details in Fig S10). (C) Rheotaxis velocity. Positive values indicate upstream movement at a flow velocity of 40 µm/s. Distribution, average, and SD of biological replicates are presented. (D) Proportion of vertical cells in rod-shaped strains. Yellow and gray colors represent vertical and horizontal cells, respectively. See *SI Appendix,* Table S6 for sample size.

Next, we examined the behavior of these 14 bacteria under a water flow of 40 µm/s and classified them into seven species and strains that exhibited positive rheotaxis and seven that did not (Fig 5C and Movie S11). All rheotactic strains and species belonged to the family *Thermaceae* with rod-shaped morphologies (Fig 5A). Among them, 5-60% of cells showed vertical orientations during water flow (Fig 5D). In contrast, five of the seven non-rheotactic species belonged to the genus *Deinococcus* with spherical or short-rod morphologies (Fig 5A), making it difficult to distinguish between vertical or horizontal orientations (Fig 5D). These results suggest a correlation between rod-shaped cell morphology and positive rheotaxis. In addition, we analyzed the flapping motion of four rheotactic species (Fig S11AB), and found a significant correlation between the upstream velocity and the apparent spring constants (Fig S11C).

## DISCUSSION

Here, we demonstrated that water flow induces upstream movement of *T. thermophilus* on surfaces (Fig 1). The optimal water flow for rheotaxis was estimated to correspond to a shear stress of 0.4 Pa (Fig 2), which is 100 times higher than that of the threshold observed for rheotaxis of *E. coli* (9). Unlike flagellated bacteria, which undergo frequentl changes of direction, *T. thermophilus* maintains a stable orientation, directing its leading pole toward water flow and sustaining unidirectional movement over a long period of time. Furthermore, T4P-dependent rheotaxis in *T. thermophilus* is 10-100 times faster than that of *P. aeruginosa* and *X. fastidiosa* (6, 28). This efficient migration may represent an evolutionary adaptation to a surface-associated lifestyle in hot spring environments with rapidly flowing water.

How does *T. thermophilus* sense external stimuli of water flow? The genome of *T. thermophilus* HB8 lacks genes involved in chemotaxis and mechanotaxis including the two-component system of Pil-Chp (29, 30) and methyl-accepting chemotaxis proteins (MCPs). This is consistent with our result that *T. thermophilus* infrequently changes its moving direction (Fig 1). In an oligotrophic environment, T4P localization is biased toward the leading pole (Fig 3 and Fig S7), suggesting that molecular mechanisms controlling T4P polarity are intrinsic to *T. thermophilus* HB8. In *Myxococcus xanthus*, T4P polarity is regulated by MglA and MglB (31), and these homologous proteins in *T. thermophilus* HB27 have been implicated in motility during colony spreading (32). However, the precise mechanism underlying monopolar-biased T4P localization in *T. thermophilus* HB8 remains unclear. Once this monopolar bias occurs, vertical cells weakly attach to the surface, allowing their cell body to tilt under shear flow (Fig 2). This tilting facilitates an upstream-biased distribution of T4P filaments at the leading pole (Fig 3). A similar strategy to weaken cell attachment during rheotaxis has been reported in *P. aeruginosa*, where the response regulator PilH increases the number of T4P filaments, resulting in a decrease of cell body attachment. (33). Considering that *T. thermophilus* lacks *pilH* homologs in the genome, this difference may reflect regulatory adaptations specific to *T. thermophilu*s, which possesses multiple T4P filaments at a single cell pole. Our results highlight the critical roles of dual retraction motors, PilT1 and PilT2, which enable rapid and frequent retraction of T4P filaments (Fig 4). This dynamic activity promotes instability in surface attachment, enhancing the sensitivity to rheotactic cues. Further studies on the subcellular localization and biochemical activity of intercellular proteins associated with rheotaxis are necessary to elucidate how *T. thermophilu*s achieves precise environmental sensing and efficient navigation.

We demonstrated that four strains of *T. thermophilu*s and three species within the family *Thermaceae* exhibit positive rheotaxis (Fig 5). The upstream-directed movement might be a common feature among *Thermaceae*, potentially enabling these bacteria to reach optimal growth environments, such as high-temperature habitats. Indeed, the shear stress in the range of 0.1-1.0 Pa, which supports the rheotactic behavior of *T. thermophilus* HB8, corresponds to the flow conditions observed in the Mine-Onsen hot spring, where this strain was originally isolated (11). Given that these values are also comparable to those measured in river environments (34), rheotaxis may be a common feature in bacteria isolated from hot springs where water flow is ubiquitous (11, 35–39). In contrast, rheotaxis was absent in two strains in *T. thermophilu*s and one species in *Thermaceae* (Table S3), which were isolated from thermophilic environments with limited water flow (40–42). From an evolutionary perspective, it is intriguing to consider why some bacteria within the same family exhibit upstream-directed movement while others do not. This variation might be explained by the distinct environmental niches in which these bacteria evolved. Conversely, all *Deinococcus* species isolated from moderate temperature environments such as soil and fish skin (19, 43–47), completely lack rheotaxis (Fig 5). A notable difference between the two groups is morphology: while *Thermaceae* bacteria are rod-shaped, *Deinococcus* species are spherical. It is plausible that the rod-shaped morphology of *Thermaceae* facilitates the detection of shear flow. This raises a new question about how *Deinococcus* utilizes surface motility and T4P dynamics for functions other than rheotaxis.

Pathogenic bacteria utilize T4P-dependent rheotaxis for efficient colonization and biofilm formation, and eventually settle in host vascular systems (28, 48, 49). By comparison, the non-pathogenic bacterium *T. thermophilus* appears to leverage water flow for long-distance migration in its natural environment. At first glance, surface motility might seem slow and inefficient compared to flagellar swimming. However, in fast-flowing environments like hot springs, this motility style likely represents an effective strategy. This highly directional migration, achieved without conventional chemotactic genes, reflects a unique adaptation to constant-flow environments. Our experimental setup, which reproduced natural water flow by controlling shear stress and temperature, sheds light on the lifestyle of surface-associated bacteria. These findings highlight how *T. thermophilus* and related species adapt to dynamic, high-temperature environments, providing broader insights into their ecological roles and contributions to the global ecosystem. Understanding such adaptations could inform future studies of microbial ecology, evolution, and the mechanisms underlying bacterial motility in diverse environments.

## Acknowledgments

The authors thank Eli Cohen for critical reading and comments on the manuscript. Strain JCM 6269, 10668, 10724, 11576, 11603, 16871, 19176, 21311, 31434, 33999, 34562, 34718 were provided by Japan Collection of Microorganisms, RIKEN BRC which is participating in the National BioResource Project of the MEXT, Japan. This study was supported by KAKENHI grants 22H05066 to DN and 24KJ1131 to NU from Japan Society for the Promotion of Science (JSPS).

## Supplemental Information

### Materials and Methods

#### Strains and culture conditions

*Thermus thermophilus* HB8 and its mutant strains were grown at 70°C in liquid NM medium (1) containing 0.8% [w/v] tryptone, 0.4% [w/v] yeast extract, 0.2% [w/v] NaCl, and 5 mM potassium phosphate (pH7.8). Bacterial species, strains, growth medium, and temperature used in this study are listed in Table S3. Cultures were grown in a petri dish with a liquid depth of about 5 mm and incubated statically. *M. rubber* and other bacterial strains were cultured to an optical density of approximately 1.0 and 0.1-0.5 at 600 nm, respectively.

#### Construction of *pilT1*, *pilT2*, and *pilT1 pilT2* mutants

All of the mutant strains were constructed by homologous recombination using integration vectors. Knockout vectors pilT1-pyrE and pilT2-pyrE (Fig S12AB) were constructed for insertional disruption of the *pilT1* and *pilT2* genes, respectively, using the *pyrE* gene, which codes for orotate phosphoribosyltransferase from the pyrimidine biosynthetic pathway. The *pyrE* gene cassette in p3TSDN1 (1) contains three termination codons for each different reading frame upstream of the Shine-Dalgarno sequence. The Δ*pyrE* strain AM114 (1) derived from T. thermophilus HB8 was transformed with these vectors and transformants were isolated in a synthetic minimum medium (2) without uracil. To construct the *pilT1 pilT2* double mutant, the *pyrE* gene inserted in the *pilT1::pyrE* strain was deleted by using the knockout vector pilFpilT1 (Fig S12C) in 5-fluoroorotic acid (5-FOA) medium as described previously (3), and then the *pyrE* gene was inserted into the *pilT2* gene using pilT2-pyrE. Oligonucleotides used in this study are listed in Table S4. More detailed methods are described below.

For insertion of the pyrE gene within the *pilT1* gene (TTHA0365), a knockout vector, pilT1-pyrE (Fig. S1a), was constructed as follows. DNA fragments encoding the N- and C-terminal regions of PilT1 were amplified by PCR using the primers sets 1A-Xho/1B-Hin and 1C-Eco/1D-Bam, respectively, with T. thermophilus HB8 genomic DNA as a template. After purification, the PCR products were digested with the restriction enzyme pairs *Xho*I/*Hind*III and *Eco*RI/*Bam*HI, respectively. The digested fragments were then sequentially cloned into the corresponding sites of the plasmid p3TSDN1, in which the *Xho*I/*Hin*dIII and *Eco*RI/*Bam*HI sites were located within the multiple cloning sites at the upstream and downstream regions of the *pyrE* gene, respectively. The resulting plasmid pilT1-pyrE was used to transform Δ*pyrE* strain AM114, and the isolated transformant in MM without uracil was designated KT204. Similarly, a knockout vector for insertion of the *pyrE* gene to inactivate the *pilT2* gene (TTHA1774), pilT2-pyrE (Fig S12B), was constructed using the two primer sets, 2A-Xho/2B-Hin and 2C-Eco/2D-Bam, and the plasmid p3TSDN1. The resulting plasmid was used to transform Δ*pyrE* strains AM114, and the isolated transformant in MM without uracil was designated KT303.

*pilT1 pilT2* double knockout strain was constructed as follows. A DNA fragment containing the C-terminal coding region of the *pilF* gene (TTHA0364) was amplified by PCR using the primers PilF-S and PilF-AS. After purification, the DNA fragment was digested with the restriction enzymes *Nde*I and *Eco*RI. The digested fragment was cloned into the corresponding sites of the plasmid pET21c. The resulting plasmid was designated pET-pilF (Fig S12C). The *Eco*RI-*Not*I fragment of pilT1-pyrE, which includes the fragment amplified with the primer set 1C-Eco/1D-Bam and encodes the C-terminal region of PilT1, was cloned at the corresponding sites of pET-pilF. The resulting plasmid pilFpilT1 contains regions encoding the C-terminus of PilF and the C-terminus of PilT1,lacking the Shine-Dalgarno sequence of the *pilT1* gene and the regions encoding the N-terminus of PilT1. pilFpilT1 was used to transform *pilT1*::*pyrE* strain KT204 to delete the *pyrE* gene inserted within the *pilT1* gene, and 5-FOA resistant clone was isolated. Then, the resulting strain KT403 from KT204, was transformed with pilT2-pyrE for insertional inactivation of the *pilT2* gene. The isolated transformant in MM without uracil was designated KT502 derived from strains KT403.

The insertion or deletion of the *pyrE* gene in the *pilT1* and/or *pilT2* gene locus was confirmed by Southern blot analysis.

### Preparation of anti-PilA antibody

For expression of a recombinant PilA in *Escherichia coli*, a fragment of the *pilA* gene (TTHA1221), spanning amino acid residues 33–122, was amplified by PCR using the primer set TTHA1221dN-S/TTHA1221dN-AS. The PCR product was digested with the restriction enzymes *Nde*I and *Eco*RI, followed by cloning into the corresponding sites of pET-28a(+). *E. coli* SHuffle T7 Express (New England Biolabs) was transformed with the resulting vector. A single colony was inoculated into 1 L of TB medium containing 25 μg/mL kanamycin. The protein was overproduced by induction with 1 mM isopropyl-beta-D-1-thiogalactoside when the optical density at 600 nm of the culture reached 0.7. After an additional 6 h at 30℃, the cells were harvested by centrifugation, washed with buffer A (20 mM Tirs-HCl, pH8.0, 150 mM NaCl), and resuspended in the same buffer. The cells were disrupted by sonication, and the cell debris was removed by centrifugation. The cell-free extract was heated at 70℃ for 10 minutes. The soluble fraction was applied to a Histrap HP (GE Healthcare), and eluted with a 35–500 mM imidazole linear gradient in buffer A. After cleavage of the N-terminal 6xHis-tag by thrombin, the recombinant protein was purified by Superdex 75 (GE Healthcare) with buffer A. Rabbit polyclonal anti-PilA antiserum was raised against the purified recombinant PilA^33-122^ protein (Hokkaido System Science, Japan).

### Optical microscopy and data analyses

Cell behavior on the glass surface was visualized under an inverted microscope (IX73 or IX83; Olympus) equipped with an objective lens (LUCPLFLN 40×PH, N.A. 0.6, UCPLFLN 20×PH, N.A. 0.7; Olympus), a CMOS camera (DMK33U174; Imaging Source), and an optical table (ASD-1510T, or HAX0605; JVI). The cell was visualized by a halogen lamp (U-LH100L-3; Olympus) through a bright-field condenser (IX2-LWUCD, NA 0.55; Olympus) for phase-contrast microscopy, or a dark-field condenser (U-DCD, NA0.8-0.92; Olympus) for dark-field microscopy. For dark-field microscopy in wide filed, the cell was visualized with a ×10 objective lens (UPLFLN 10×2PH; N. A. 0.3; Olympus) and LED ring light (TLED-JZ02; Tledtech) used as a dark-field condenser. The microscope stage was heated with a thermoplate (TP-110R-100; Tokai Hit). Projections of the images were acquired with an imaging software IC Capture (Imaging Source) under 1-s resolution and converted into an AVI file without any compression. All data were analyzed by Fiji (ImageJ 1.53t) (4) and its plugin, TrackMate (5). If necessary, the drift of the images was corrected by the plugin Manual Drift Correction, and the luminance unevenness of the images was processed by Subtract Background.

For immunofluorescence microscopy of T4P filaments, the sample was examined under the inverted microscope equipped with a ×100 objective lens (UPLXAPO100XOPH, N.A. 1.45; Olympus), a filter set (Cy3-4040C, Semrock), the CMOS camera, and the optical table. The fluorescent signal of T4P filaments was visualized by a mercury lamp (U-HGLGPS; Olympus) or collimated green-light LED (M530L4, Thorlabs). The projection of the image was captured with the imaging software IC Capture and converted into a TIFF file without any compression.

For the bead assay, the cells and microbeads were visualized under the inverted microscope equipped with the ×40 objective lens, a filter set (GFP-4050B, Semrock), the CMOS camera, and the optical table. The position of the cell and microbeads was visualized by a halogen lamp through a dark-field condenser and a collimated blue-light LED (M470L5; Thorlabs). The microscope stage was heated with the thermoplate. Projections of the images were acquired with the imaging software IC Capture under 0.1-s resolution and converted into an AVI file without any compression.

### Cell preparation for twitching motility

The motility buffer used in this study was composed of 0-500 mM NaCl and 5 mM potassium phosphate at a pH adjusted for each strain as listed in Table S5. For observation of the genus *Thermus*, *Marinithermus*, and *Meiothermus*, the cell culture was centrifuged at 12,000 × g for 2 min, and the pellet was suspended in the motility buffer at the same volume as the original culture. The suspension was subsequently used for the observation for flow experiment, bacterial motility, and bead’s assay. For immunofluorescence microscopy of PilA, the cell was suspended in the motility buffer at one-tenth of the original culture volume. For observation of the genus *Deinococcus*, the cell culture was used for the flow experiment directly and washed with the motility buffer in the flow chamber as described in the flow experiment section.

### Construction of the flow chamber

The flow chamber was assembled by taping a coverslip with a glass slide, as described previously (6). The glass slide was bored with a high-speed drill press equipped with a diamond-tipped bit (1 mm diameter, No. 13853, NAKANISHI) for inlet and outlet ports. The center portion of the double-sided tape (665-3-24; 3M or 7082; Teraoka) was cut out 25 mm long and 1.5 mm wide. The chamber was assembled by taping the coverslip with the glass slide. Inlet and outlet ports (N-333; IDEX Health & Science) were attached with high-temperature elastic adhesive (Super X No. 8008 clear; CEMEDINE). The finished channel of the sample chamber was straight with the dimensions of width: 1.5 mm, height: 0.1 mm, and length: 25 mm for bright-filed microscopy, or 50 mm for dark-field microscopy. A syringe pump (Legato 200; Kd Scientific) was connected to the flow chamber by a connecter and tube (F-333NX and 1512L; IDEX Health & Science). The flow chamber and the tube coiled for a length of 256 mm were heated on the thermoplate on the microscope stage. The temperature of the chamber and thermoplate on microscope stage was measured by thermography (C3-X; FLIR).

### Flow experiments under temperature control

Cell suspensions were injected into the chamber and incubated for 4 min. For the observation of cells belong the genus *Thermus, Marinithermus,* and *Meiothermus*, the flow chamber was pre-coated with BSA buffer containing 2% BSA (wt/vol) in the motility buffer, and then washed with motility buffer. The motility buffer was degassed by an aspirator (AS-01; AS ONE) for 2 hours. The unattached cells were slowly washed away by the motility buffer for nutrient-free conditions and by the flesh medium for nutrient-rich conditions, respectively, at a flow rate of 1 µL/s for 1 min. The behavior of the attached cells in fluid flow was observed. Each flow rate of the syringe pump was calibrated to the velocity of the fluid flow near the glass surface at 70℃. The velocity was determined from the flowing cell that moved passively in the flow direction immediately after the occasional detachment from the glass surface. The shear stress at the surface of the chamber was estimated with the equation described previously (7).

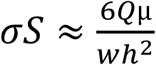

Where *Q* is the flow rate; *µ* is the viscosity of the solution, which is assumed equal to that of water; *w* and *h* are the width and height of the flow chamber, respectively.

### Electron microscopy

Samples bound to the grids were stained with 2% (wt/vol) ammonium molybdate and observed by transmission electron microscopy, as previously described (8). Carbon-coated EM grids were glow-discharged by a PIB-10 hydrophilic treatment device (Vacuum Device) before use. Bacterial culture was put on the EM grid and incubated for 3 min at 65℃. The cells were chemically fixed with 1% (vol/vol) glutaraldehyde in the motility buffer for 10 min at RT. After washing three times with the buffer, the cells were stained with 2% ammonium molybdate and air-dried. Samples were observed under a transmission electron microscope (JEM-1400, JEOL) at 100 kV. The EM images were captured by a charge-coupled device (CCD) camera and analyzed by ImageJ 1.48v.

### Measurement of twitching motility in horizontal cells without fluid flow

A tunnel chamber was assembled by taping coverslips with double-sided tape (∼90 μm thick, NW-5; Nichiban) (9). The chamber was pre-coated with BSA buffer and washed with motility buffer. Cell suspensions of *T. thermophilus* HB8 WT and the T4P mutant were poured into the chamber and incubated at 70℃ for 4 min on the stage with the thermoplate. The unattached cells were washed away with motility buffer, and then the ends of the chamber were sealed with nail polish.

### Correlative immunofluorescence microscopy of T4P filaments

The coverslip coated with carbon through a locator grid (MAXTA FORM H2; Nisshin EM) was prepared by evaporation in a vacuum evaporator (VE-2012, Vacuum Device). The coverslip coated by thin-carbon with the grid pattern was used for the cell positioning. The tunnel chamber assembled by taping the patterned coverslip with double-sided tape was pre-coated with the BSA buffer and washed with the motility buffer. The cell suspension was poured into the tunnel chamber and incubated at 70℃ for 4 min. The unattached cells were washed away by the motility buffer. The behavior of the attached cells was captured under 1-s resolution before and during the fixation. Cells were chemically fixed with 1% (vol/vol) glutaraldehyde in the motility buffer for 5 min at 70℃. After washing with the motility buffer, the cells were incubated in the BSA buffer for 5 min at room temperature (RT). Cells were incubated with the primary antibody anti-PilA in the BSA buffer for 15 min at RT, followed by washing three times with the motility buffer. Cells were incubated with the secondary antibody goat anti-rabbit, Cy3 conjugate (Sigma-Aldrich) in the BSA buffer for 15 min at RT followed by washing three times with the motility buffer. Both ends of the chamber were sealed with nail polish, and the remaining cells on the glass surface were observed under fluorescent microscopy. The cell behaviors before fixation and the localization of T4P filaments by fluorescent microscopy were correlated with the grid pattern on the coverslip.

### Bead assay for visualizing T4P dynamics

Fluorescent silica beads at a size of 0.2 µm in diameter (sicastar®-greenF; micro mod, 50 mg/ml) were diluted 1/20 in the BSA buffer followed by washing three times with the motility buffer. The beads were diluted to a final concentration of 0.5 mg/ml in the motility buffer. A coverslip was glow-discharged by a hydrophilic treatment device (YHS-R; SAKIGAGE). The cell suspension was poured into a tunnel chamber assembled by taping hydrophilic coverslips. After incubation at 70℃ for 4 min, the diluted fluorescent beads were added to the chamber and washed away the unattached cells. Both ends of the chamber were sealed with nail polish, and the chamber was used for the observation.

### Mathematical simulation

All the code and data to reproduce the numerical results reported in this manuscript are available from the authors upon reasonable request.

## Supplemental Text

### Mathematical model of bacterium moving on a planar surface using T4P filaments

In order to reproduce the twitching motility of *Thermus thermophilus* on the planar surface, we propose an individual model in which the cell body is regarded as a prolate spheroid with a major radius *a* and a short radius *b* (Fig S13A). The T4P filament is regarded as a linear spring with a spring constant *k*, and the extension and retraction of the T4P filaments due to the polymerization and depolymerization of pilin are modeled by changing the equilibrium length of the spring.

When a T4P filament is extended without its tip adsorbed on the planar surface, the equilibrium length of the spring increases at a constant rate *V*_*e*_ (Fig S13B). In this case, the T4P filament extends linearly in any direction within a cone with the half apex angle *β* whose axis of rotation is normal to the planar surface tangent to the base of the filament. If the tip of the T4P filament contacts the planar surface, the T4P filament attached to the planar surface and turns from extension to retraction. If the equilibrium length of the spring exceeds the maximum value *d*_*max*_ before it contacts the planar surface, the filament extension will also change to retraction. The retraction is achieved by shortening the equilibrium length of the spring at a constant rate *V*_*r*_. If *t*_*i*_ = 0 is the time when the tip of T4P filament *i* is attached to the planar surface, the tension ***F***_*r*,*i*_(*t*_*i*_) at time *t*_*i*_ is expressed as

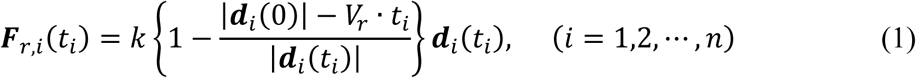

where ***d***_*i*_(*t*_*i*_) is the direction vector from the base of the T4P filament *i* to the attachment point at time *t*_*i*_ . If the magnitude of the tension |***F***_*r*,*i*_(*t*_*i*_)| is greater than *F*_*max*_, the tip of the T4P filament is detached from the planar surface, and the tension then becomes zero. In addition, when the condition of |***d***_*i*_(0)| ≦ *V*_*r*_ · *t* is satisfied, the tip of the T4P filament is detached from the planar surface, and the retracted filament is changed to be extended. By repeating the retraction and extension of the T4P filament with the attachment and detachment of the tip to the planar surface, the cell body translates and rotates in three-dimensional space while performing twitching motility.

Since the tips of multiple T4P filaments repeatedly attach to and detach from the planar surface asynchronously, the probability that the cell body will move far away from the planar surface is low. However, in order to avoid the cell body from slipping through the planar surface, potential energy *U*_LJ_(*h*, *θ*) is introduced between the cell body and the planar surface (Fig S13C).

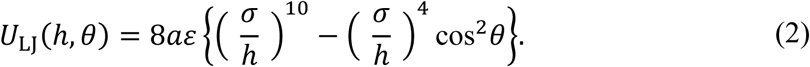

Equation (2) is used as the interaction potential between the liquid crystal molecules and the substrate (10), and both attractive and repulsive forces also act between the cell body and the planar surface. Here, *h* is the minimum distance between the planar surface and the cell body, *θ* is the angle between the major axis vector of the cell body and the planar surface, *ɛ* is the depth of the binding potential, and *σ* is the distance *h* when the potential energy becomes zero. The hydrodynamic interaction between the cell body and the planar surface is not considered to simplify the model.

The force on the prolate spheroid, which is considered as the cell body, is calculated by the sum of the tension of *n* T4P filaments 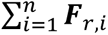 and the partial derivative of the potential energy −**▽***U*_LJ_(*h*, *θ*) from Equation (2). If the combined force is ***F*** , the torque around the principal axis of inertia of the prolate spheroid is ***T̃***_*P*_ , and the viscosity coefficient of the fluid in the tail around the cell body is *η* , the velocity ***v*** of the hydrodynamic center ***r*** = (*x*, *y z*) of the cell body and the angular velocity ***ω̃*** _*P*_ of the rotation around the principal axis of inertia are expressed as follows from Stokes’ resistance law (11).

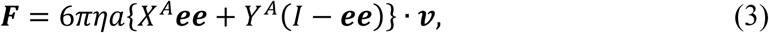

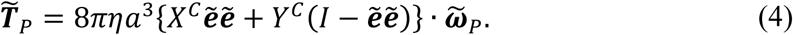

where ***e*** is the unit vector in the direction of the major axis of the cell body, and the direction of the head is positive. Variables with tildes such as ***ẽ***, ***T̃***_*P*_ and ***ω̃***_*P*_ represent vectors whose coordinate axis is the principal axis of inertia of the cell body. That is, ***ẽ*** = (1, 0, 0). ***ee*** and ***ẽẽ*** are the binomial products of ***e*** and ***ẽ***, respectively, and *I* is the unit tensor. *X*^*A*^ , *Y*^*A*^ , *X*^*C*^ and *Y*^*C*^ are the resistance tensor components of a prolate spheroid mimicking the cell body, and they are expressed as follows (11):

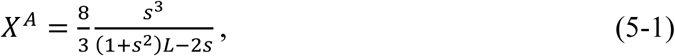

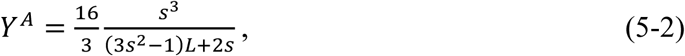

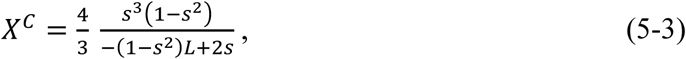

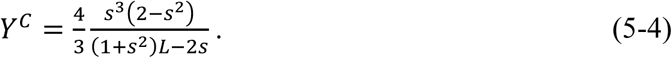

Here, *L* = ln{(1 + *s*)⁄1 − *s*} and 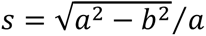. Equations (3) and (4) can be solved using the Runge-Kutta method with a time step of Δ*t* = 0.001 to calculate the time variation of the center coordinate ***r*** and the unit vector ***e*** of the cell body. Note that it is not appropriate to calculate fluid flow due to external force fields.

**Fig S1.**
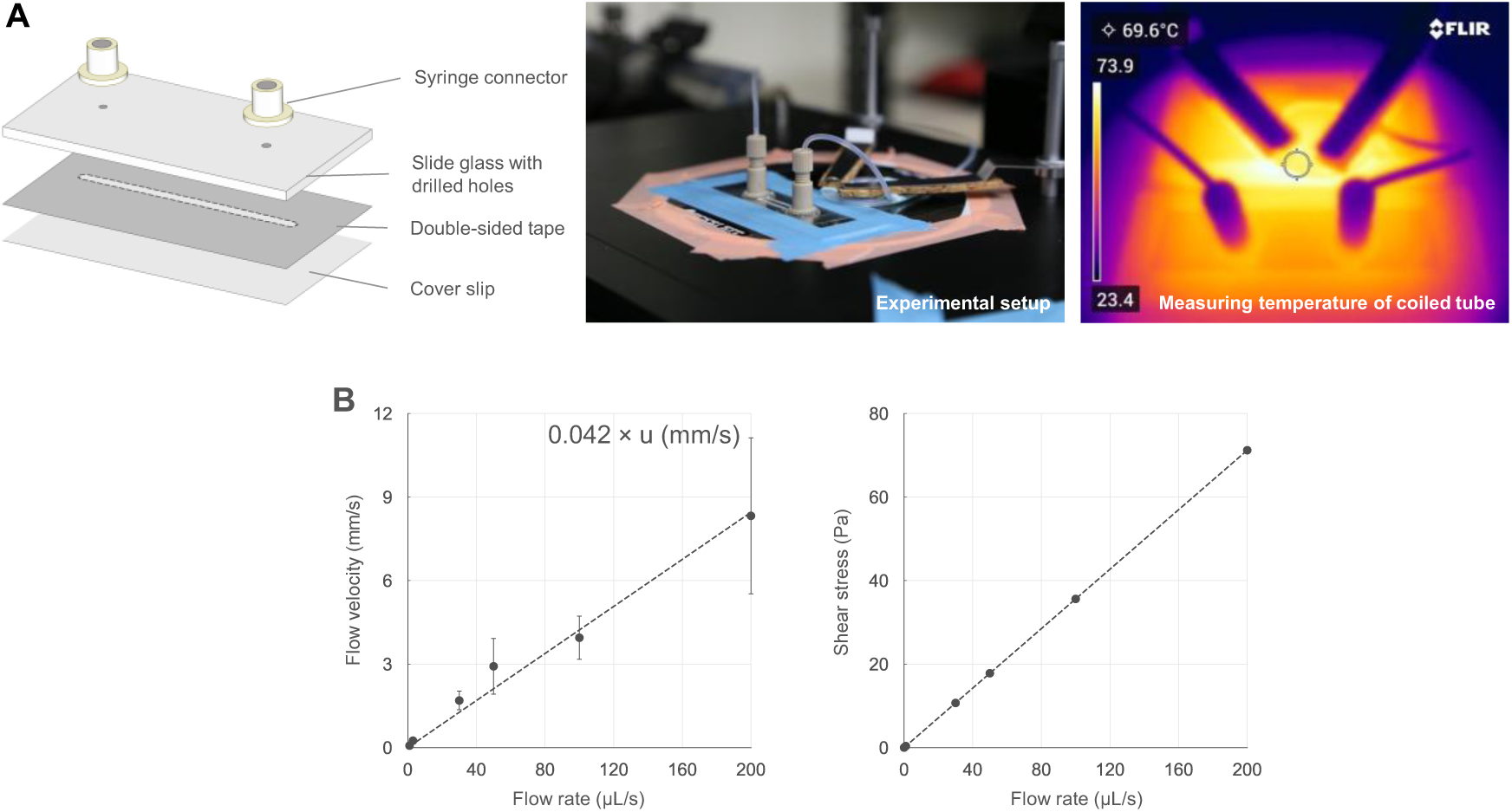
Temperature-controlled flow experiment. (A) Experimental setup. *Left*: Schematic of the flow chamber assembly. *Middle*: Image of the flow chamber and tube heated on a stage heater. *Right*: Thermographic image, showing temperature distribution. The center circle temperature is presented in the upper left corner. (B) Calibration of flow dynamics. *Left*: Relationship between flow rate of syringe pump and flow velocity near the glass surface, measured from the initial velocity of detached cells (n = 3-14 cells at each data point). *Right*: Relationship between flow rate of syringe pump and shear stress calculated based on the geometry of flow chamber.

**Fig S2.**
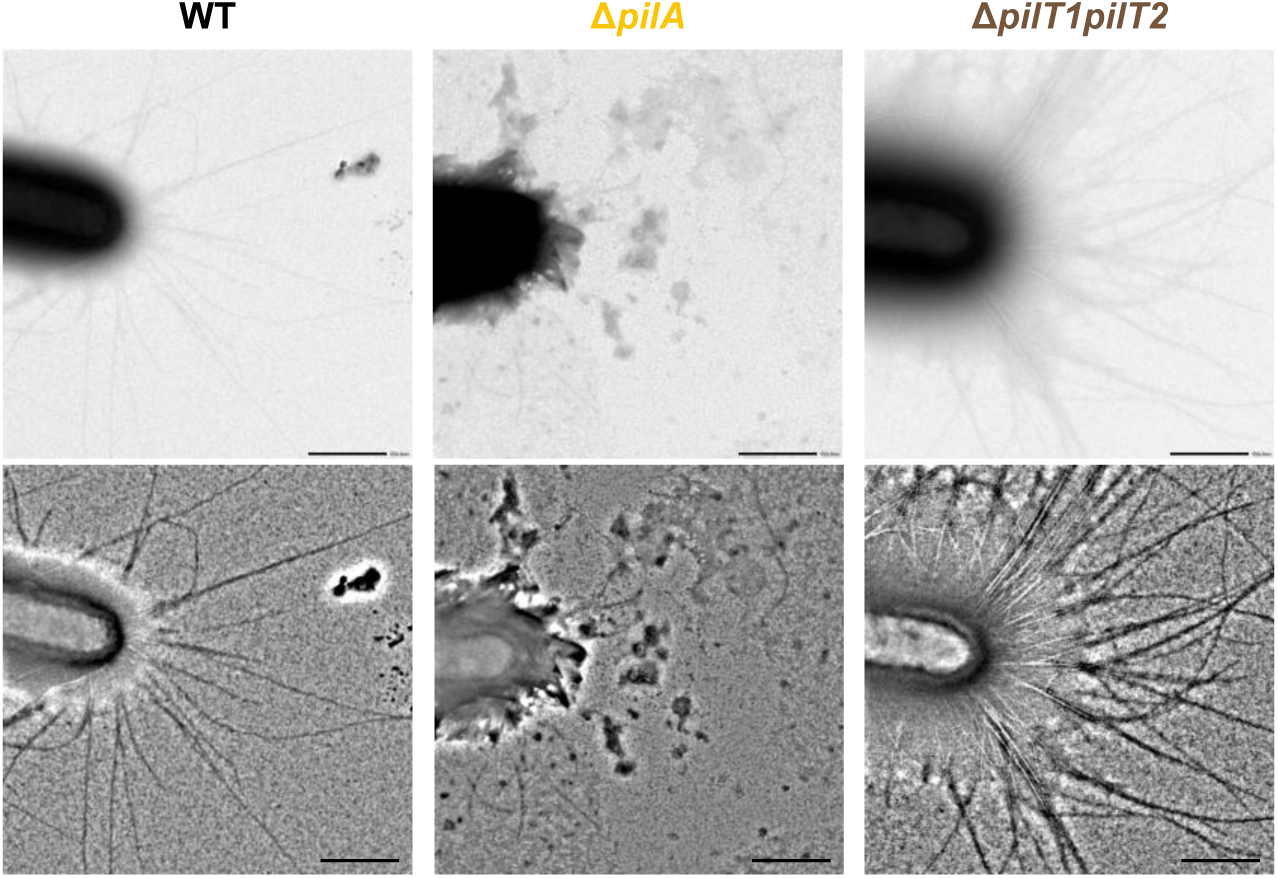
Negative staining EM images of cells of *T. thermophilus* HB8 WT, Δ*pilA*, and Δ*pilT1pilT2* mutants. Upper images: Originals. Lower images: Bandpass-filtered images to enhance the surface filaments. Scale bar, 500 nm.

**Fig S3.**
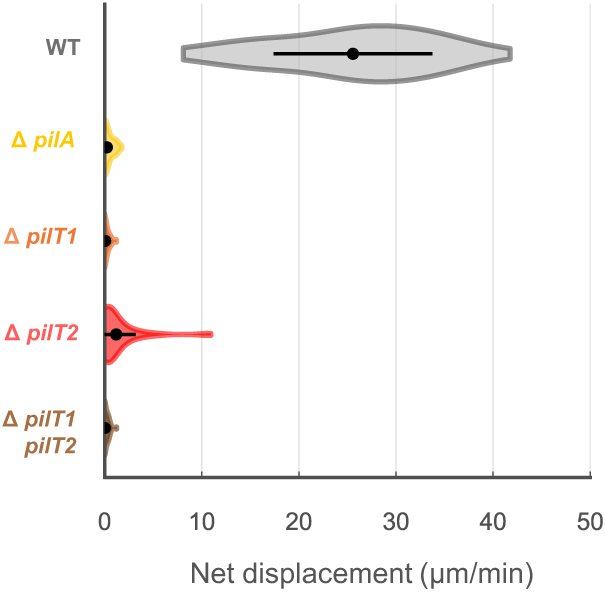
Surface movement of horizontal cells. Net displacement of horizontal cells in *T. thermophilus* HB8 WT and T4P mutants for 1 min under the nutrient-free condition without water flow. The distance of surface movement was measured from its initial to final position. Distribution, average, and SD of biological replicates are presented. n = 41 (WT), 56 (Δ*pilA*), 41 (Δ*pilT1*), 75 (Δ*pilT2*), 63 (Δ*pilT1pilT2*).

**Fig S4.**
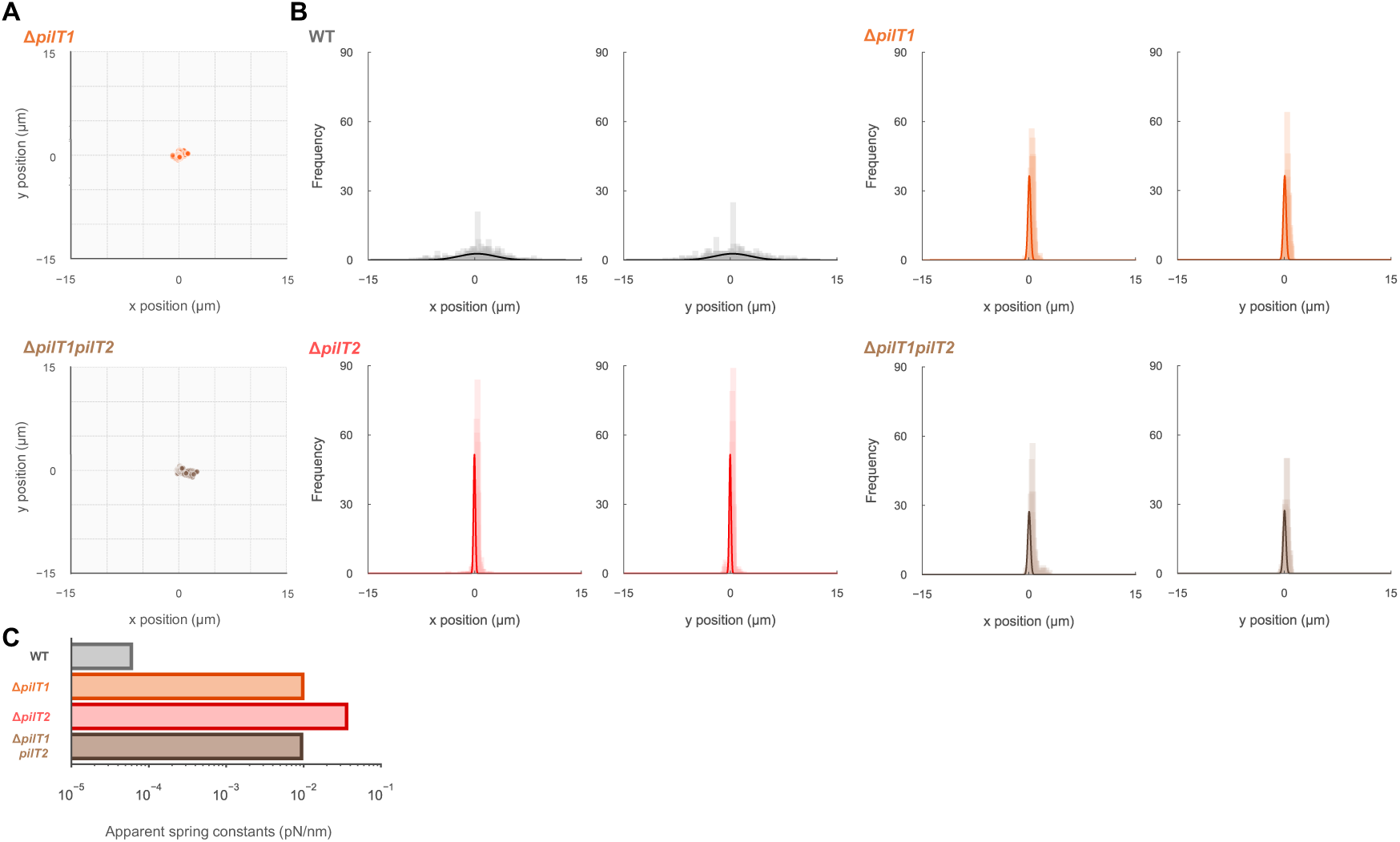
Flapping motion of vertical cells. (A) Distribution of unattached pole position relative to its attached pole during surface movement of Δ*pilT1* and Δ*pilT1pilT2* mutant cells without water flow (n = 450 from 15 cells at 1-s intervals for 30 s). (B) Distribution of unattached pole position derived from the same datasets as panel A and Fig 2E. The solid line shows the Gaussian fitting. (C) Apparent spring constants estimated from the Gaussian fitting of y positions in panel B.

**Fig S5.**
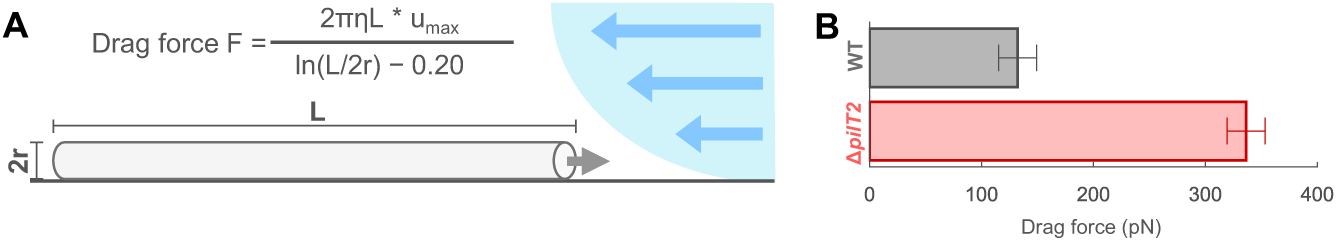
Maximum drag force in *T. thermophilus* HB8 WT and Δ*pilT2*. (A) Schematic illustration of drag force calculation. The drag coefficient was given by the cell shape assuming as a cylindrical rod with a length of 7 µm and a diameter of 0.5 µm. (B) Maximum drag force. Drag force was estimated from the flow velocity, where all vertical cells detached from the surface. Average and SD of technical replicates are presented (n = 3 for each strain). Calibration of high flow velocity was performed separately from low flow rates.

**Fig S6.**
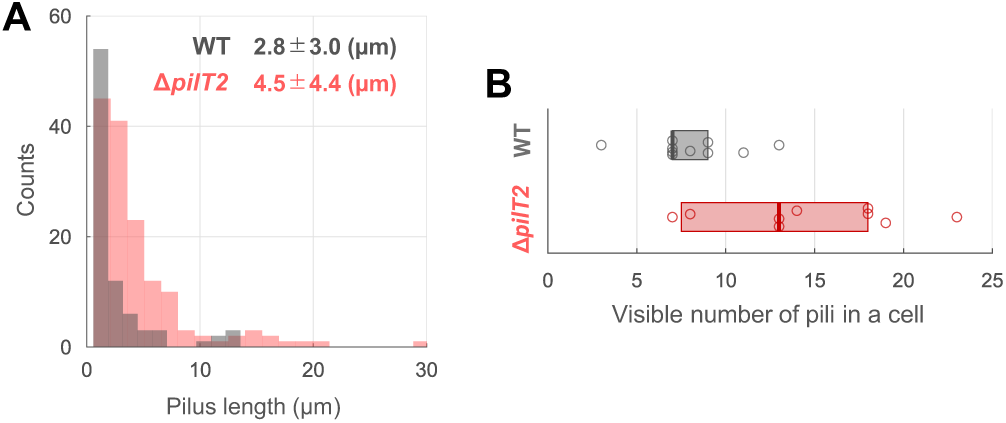
Length and number of T4P filaments in vertical cells. (A) Length distribution of T4P filament in vertical cell of *T. thermophilus* HB8 WT and Δ*pilT2* (n = 84 and 148 from 11 cells of WT and Δ*pilT2*). (B) Visible number of T4P filaments in single vertical cells of WT and Δ*pilT2* (n = 11 cells for each strain). Circles indicate biological replicates, and boxplots represent the median and 25%/75% quantile.

**Fig S7.**
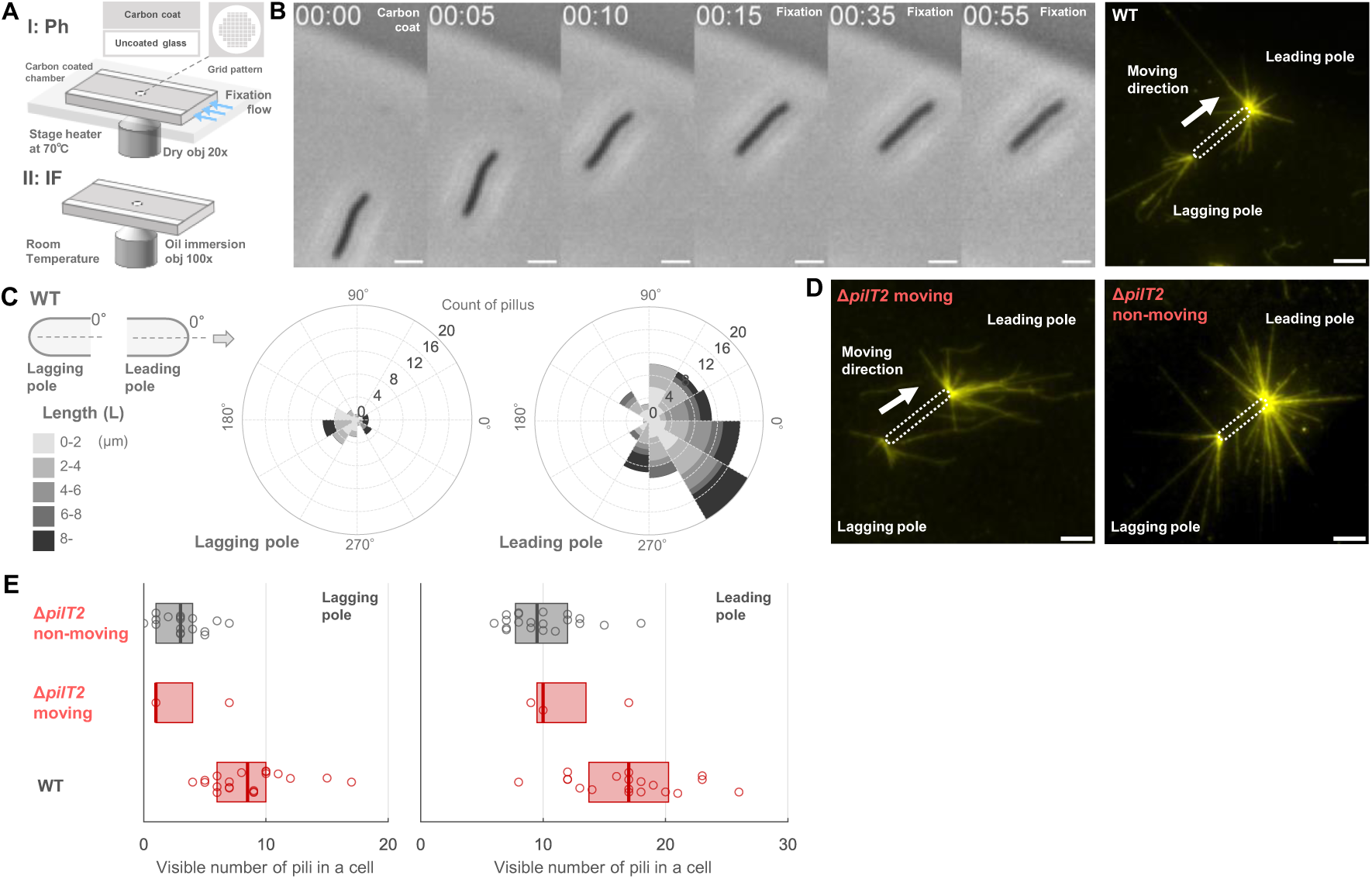
Visualization of T4P filaments in horizontal cells. (A) Schematic diagram of correlation microscopy. I: Observation of cell behavior at 70°C on a stage heater under phase microscopy using a dry objective lens (low magnification). II: Immunofluorescence microscopy at RT using oil immersion objective lens (high magnification). Carbon-coated coverslips with a grid pattern were used in tunnel chambers (see more details in *SI Appendix*, Materials and Methods section). (B) Correlation of cell movement and T4P filaments localization. Left: Time-lapse phase-contrast images under water flow applied from the right. WT cells were chemically fixed under the nutrient-free condition at the time point of 15 s. Right: Immunofluorescence image of PilA. White dashed lines outline the cell body. Scale bar, 3 µm. (C) Length and distribution of T4P filaments. Left: Schematic of the measured parameters. θ: angle between the filament and the cell body, where the moving direction corresponds to angle zero. L: length of the T4P filaments. Right: Rose plot showing angle distribution and length of T4P filaments at the lagging (middle) and leading (right) poles (n = 11 cells). (D) Immunofluorescence image of PilA in horizontal Δ*pilT2* cells. A moving (left) and non-moving cell (right). (E) Number of visible T4P filaments in single horizontal cells. Comparisons between leading (right) and lagging poles (left) of WT (top, n = 20), moving Δ*pilT2* (middle, n = 3), and non-moving Δ*pilT2* cells (bottom, n = 20). Circles indicate biological replicates, and boxplots represent the median and 25%/75% quantile.

**Fig S8.**
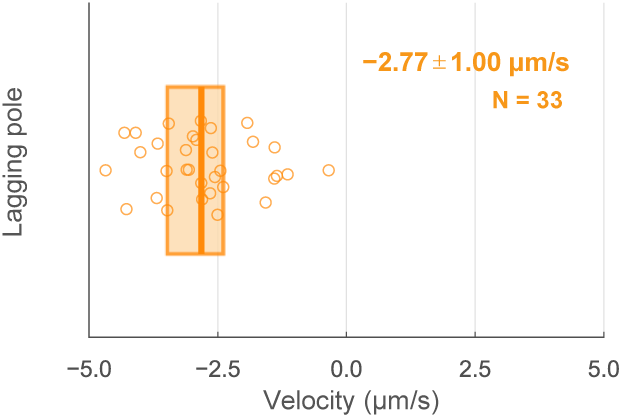
Visualization of T4P dynamics in the lagging cell pole. The velocity of bead movement towards the lagging pole of *T. thermophilus* HB8 WT cells. The velocity was determined by linear fitting of the bead displacement. Directional bead movements for more than 0.5 s were used for data analyses. Bead movement towards the cell pole is defined as a negative value. Circles indicate biological replicates, and a boxplot represents the median and 25%/75% quantile (n = 9 cells).

**Fig S9.**
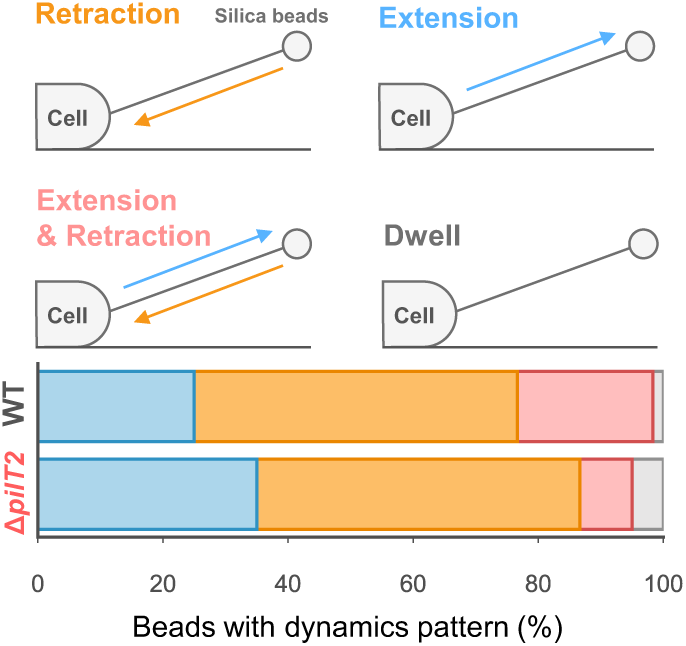
Fraction of the bead dynamics. Bead movements at the leading pole of horizontal cells in *T. thermophilus* HB8 are classified into four groups presented at the top (n = 60 events for each strain and n = 4, 12 cells for WT and Δ*pilT2*).

**Fig S10.**
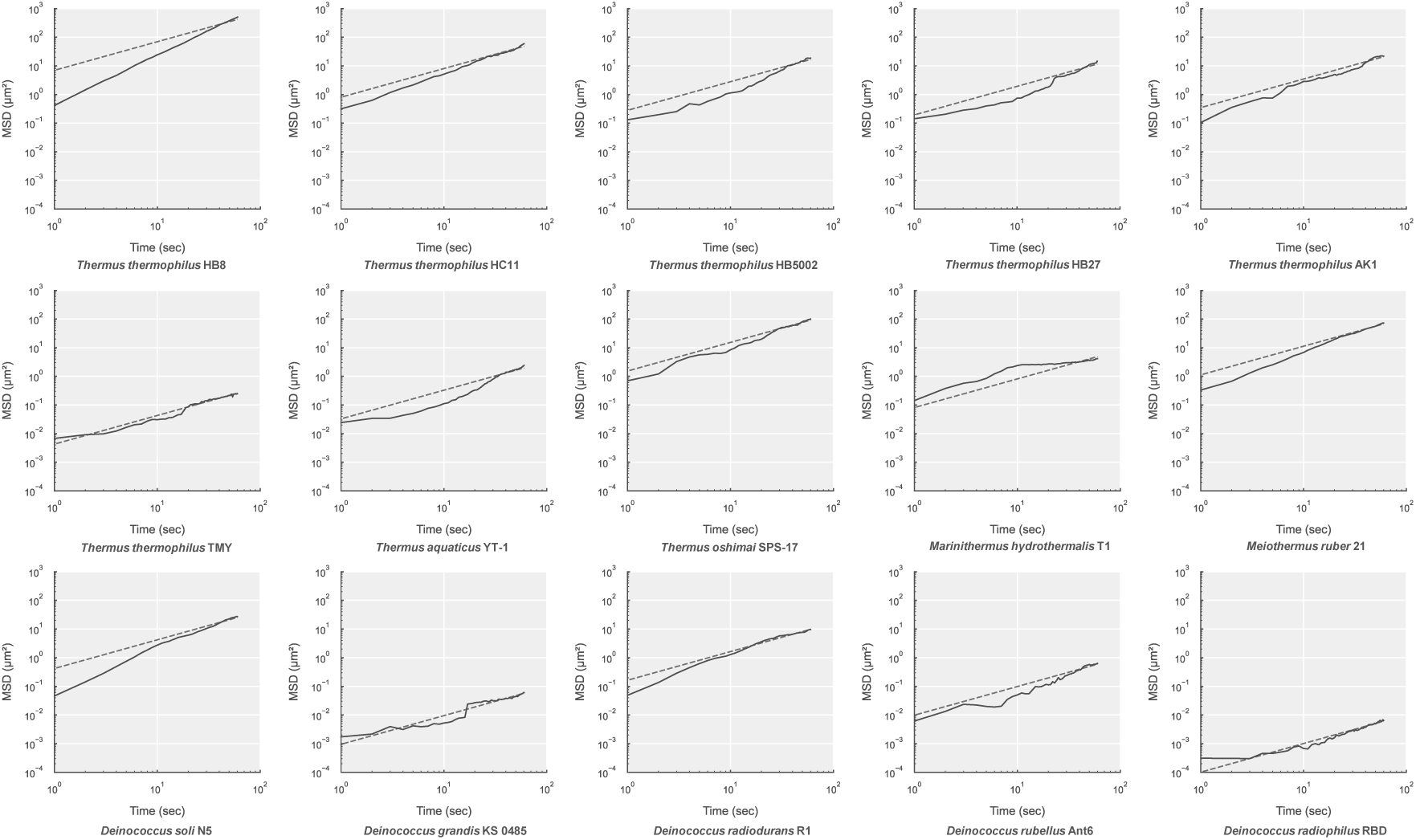
Twitching motility in *Deinococcus-Thermus*. Mean square displacement (MSD) plots of cell movement in each strain. Cells were observed under the condition without water flow. Horizontal cells are used for the data analysis in *Thermus*. Solid and dashed lines show the average and the linear fitting of 60 s, respectively. See *SI Appendix,* Table S6 for sample sizes.

**Fig S11.**
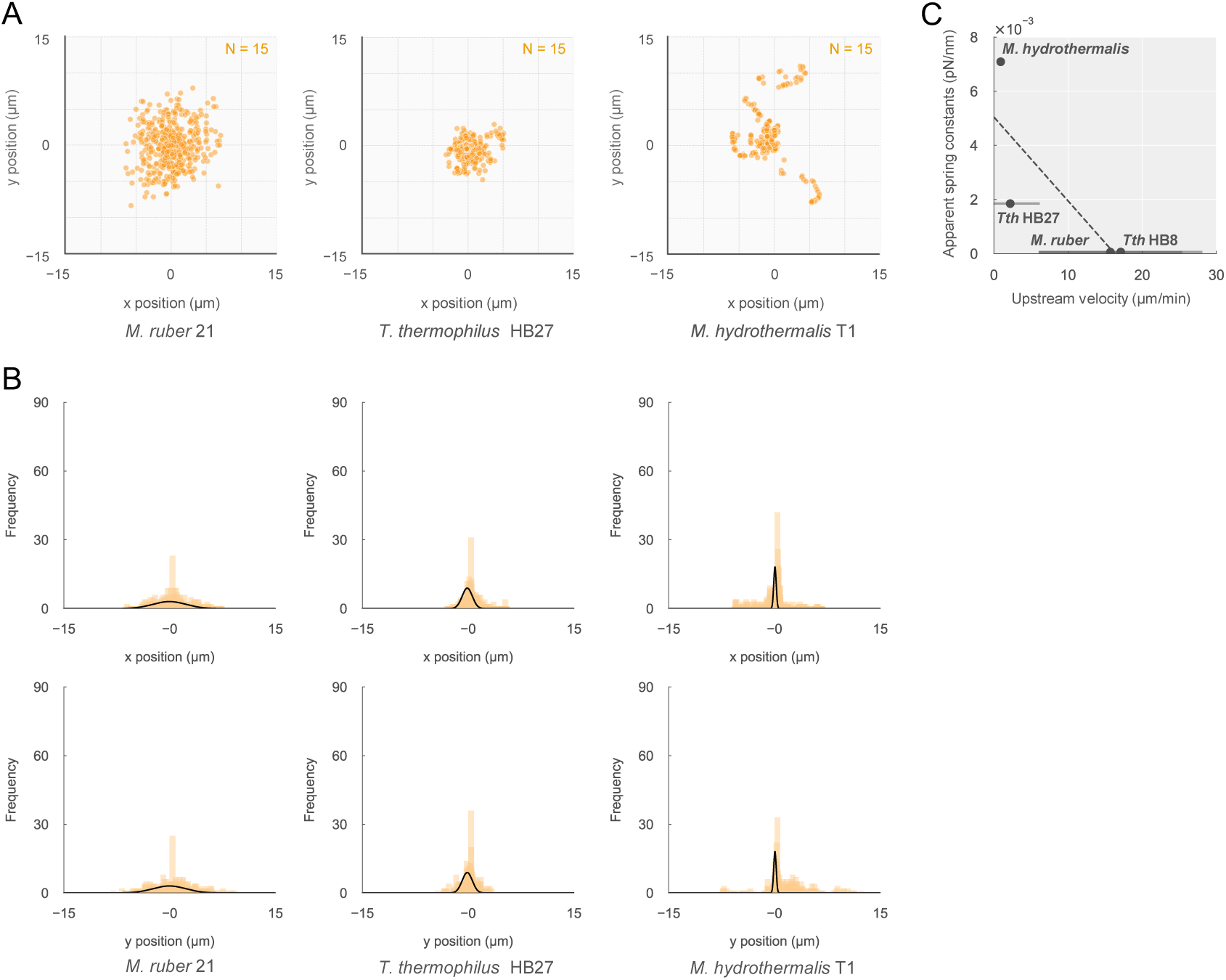
Flapping motion in *Deinococcus-Thermus*. (A) Distribution of unattached cell pole position relative to attached cell pole (n = 450 from 15 cells at 1-s intervals for 30 s). Vertical cells of *M. ruber* 21 (left), *T. thermophilus* HB27 (middle), and *M. hydrothermalis* (right) were analyzed under the condition without water flow. (B) Distribution of unattached cell pole position along the x and y axes from the data in panel A. The black line shows the Gaussian fitting. (C) Apparent spring constants from panel B. The data set of *T. thermophilus* HB8 derived from the same experiments in Fig S4. (D) Relationship between the apparent spring constant and the velocity of rheotaxis. The spring constants were derived from the y-axis data in panel C, while velocities of rheotaxis were data from Fig 5C. Error bars show SD of biological replicates in the velocity of rheotaxis.

**Fig. S12.**
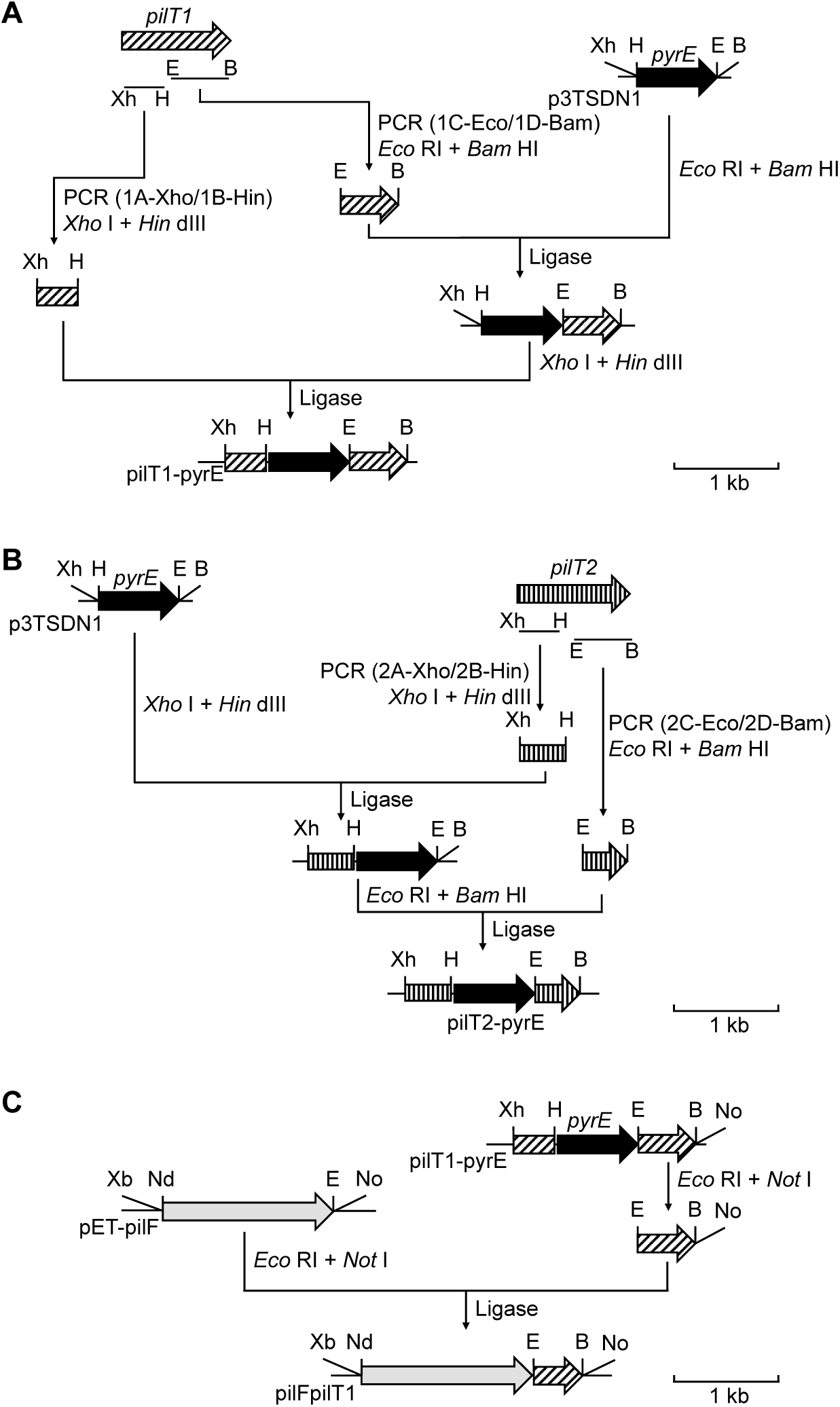
Construction of the integration vectors. (A) pilT1-pyrE for inactivation of the *pilT1* gene to construct the *pilT1* strain KT204. (B) pilT2-pyrE for inactivation of the *pilT2* gene to construct the *pilT2* strain KT303, and the *pilT1 pilT2* double mutant KT502. (C) pilFpilT1 for deletion of the *pyrE* gene of KT204.

**Fig. S13.**
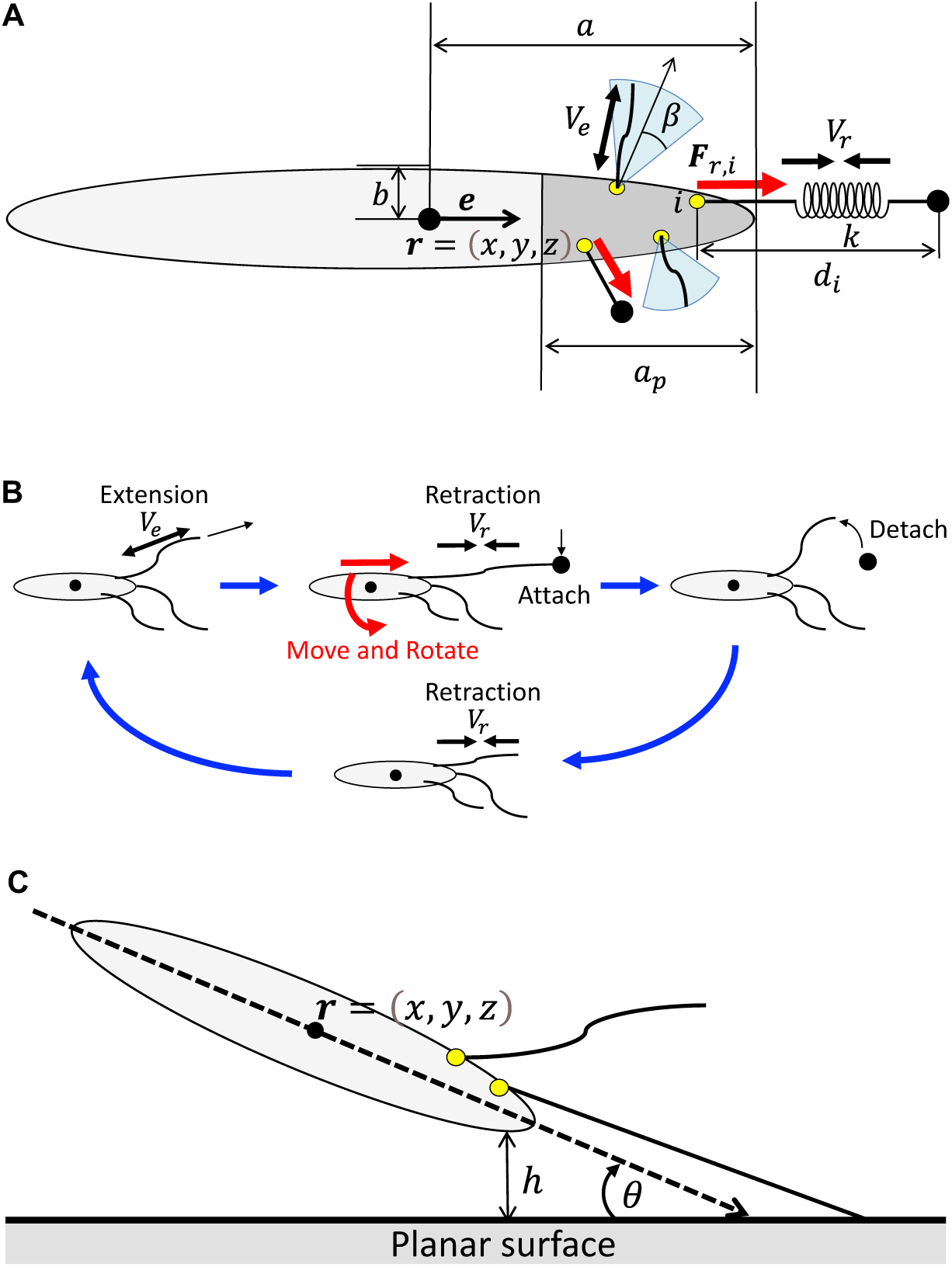
Mathematical model of a bacterium moving with T4P filaments. (A) A prolate spheroid is considered as a cell body. T4P filaments grow in the region of length *a*_*p*_ from the tip of the head to the tail. (B) The diagram of the cycle of T4P filament extension → tip attachment → retraction → tip detachment → retraction → extension. (C) The diagram of the interaction between the cell body and the planar surface.

**Table S1.** Parameters used in mathematical simulation.

| Strain | Number of<br>pili<sup>a</sup> $n$ | Retraction force<br>limit<sup>b</sup> $F_{max}$ | Retraction<br>speed<sup>c</sup> $V_r$ | Extension<br>speed<sup>c</sup> $V_e$ | Pili distribution<br>from tip of cell<sup>d</sup> $a_p$ |
| --- | --- | --- | --- | --- | --- |
| HB8 WT | 8 | 16.5 pN | -3.04 $\mu\text{m/s}$ | 1.01 $\mu\text{m/s}$ | 0.32 $\mu\text{m}$ |
| HB8 $\Delta pilT2$ | 14 | 24.1 pN | -1.27 $\mu\text{m/s}$ | 0.65 $\mu\text{m/s}$ | 0.42 $\mu\text{m}$ |

<sup>a</sup>Number of pili was measured in Fig S6B.
<sup>b</sup>Retraction force limits were estimated from drag force (Fig S5B) divided by the number of pili.
<sup>c</sup>Retraction and extension speed were measured in Fig 4D.
<sup>d</sup>The distance from the tip of cell body where the pili are distributed. The distance $0.32 \pm 0.10$ $\mu\text{m}$ in WT and $0.42 \pm 0.12$ $\mu\text{m}$ in $\Delta pilT2$ were measured at the leading pole of horizontal cells. (n = 12 and 10 cells for WT and $\Delta pilT2$ )

**Table S2.** Basic parameters used in mathematical simulation.

| Cell<br>length<br>$2a$ | Cell<br>diameter<br>$2b$ | Spring<br>constant<br>of pili $k$ | The half apex angle of<br>the cone in which the<br>pili can fluctuate $\beta$ | Max length<br>of pili<sup>a</sup> $d_{max}$ | Viscosity $\eta$ | Adhesion force<br>of cell body $\varepsilon$ | Distance between cell<br>body and planar surface<br>with the potential<br>energy zero $\sigma$ |
| --- | --- | --- | --- | --- | --- | --- | --- |
| 7.0 $\mu\text{m}$ | 0.5 $\mu\text{m}$ | 0.5 pN/$\mu\text{m}$ | 30° | 30 $\mu\text{m}$ | 0.0004 Pa·s | 0.04 pN | 0.5 $\mu\text{m}$ |

<sup>a</sup>Maximum length of pili was 30 $\mu\text{m}$ in Fig S6A.
<sup>b</sup>Viscosity of water at 70°C.

**Table S3.** Strains and culture conditions.

| Strain | Medium | Temp (°C) | Reference/Source | Isolated place |
| --- | --- | --- | --- | --- |
| *Thermus thermophilus* HB8<sup>T</sup> WT | NM | 70 | Tamakoshi Lab (1) | Mine-Onsen<br>hot spring |
| *Thermus thermophilus* HB8 $\Delta pilA$ | NM | 70 | Tamakoshi Lab (1) | - |
| *Thermus thermophilus* HB8 $\Delta pilT1$ | NM | 70 | This study | - |
| *Thermus thermophilus* HB8 $\Delta pilT2$ | NM | 70 | This study | - |
| *Thermus thermophilus* HB8 $\Delta pilT12$ | NM | 70 | This study | - |
| *Thermus thermophilus* HB27 | NM | 70 | Tamakoshi Lab (1) | Mine-Onsen<br>hot spring |
| *Thermus thermophilus* HB5002 | TM BROTH<br>JCM 273 | 70 | JCM 34562 (12) | Mine-Onsen<br>hot spring |
| *Thermus thermophilus* HC11 | TM BROTH | 70 | JCM 33999 (13) | Mine-Onsen<br>hot spring |
| *Thermus thermophilus* AK1 | TM BROTH | 70 | JCM 34718 (14) | Arima hot spring |
| *Thermus thermophilus* TMY | TM BROTH | 75 | JCM 10668 (15) | Silica scale of<br>geothermal plant |
| *Thermus aquaticus* YT-1<sup>T</sup> | CASTENHOLZ<br>JCM 276 | 70 | JCM 10724 (16) | Hot spring in Yellow<br>Stone National Park |
| *Thermus oshimai* SPS-17<sup>T</sup> | CASTENHOLZ | 65 | JCM 11603 (17) | Hot spring in Sao<br>Pedro do Sul |
| *Marinithermus hydrothermalis* T1<sup>T</sup> | Marine broth 2216<br>JCM41 | 70 | JCM 11576 (18) | Deep-sea<br>hydrothermal vent<br>chimney |
| *Meiothermus ruber* 21<sup>T</sup> | *Thermus*<br>NBRC 1104 | 55 | NBRC 106122 (19) | Hot spring in<br>Kamchatka<br>peninsula |
| *Deinococcus soli* N5<sup>T</sup> | R2A<br>JCM 346 | 30 | JCM 19176 (20) | Rice field soil |
| *Deinococcus grandis* KS0485<sup>T</sup> | NUTRIENT No2<br>JCM22 | 30 | JCM 6269 (21) | Eurasian carp<br>intestines |
| *Deinococcus radiodurans* R1<sup>T</sup> | NUTRIENT<br>JCM663 | 30 | JCM 16871 (22) | Irradiated beef can |
| *Deinococcus rubellus* Ant6<sup>T</sup> | R2A | 25 | JCM 31434 (23) | Antarctic fish<br>muscle |
| *Deinococcus radiophilus* RBD<sup>T</sup> | NUTRIENT | 30 | JCM 21311 (24) | Irradiated Bombay<br>duck |

**Table S4.**
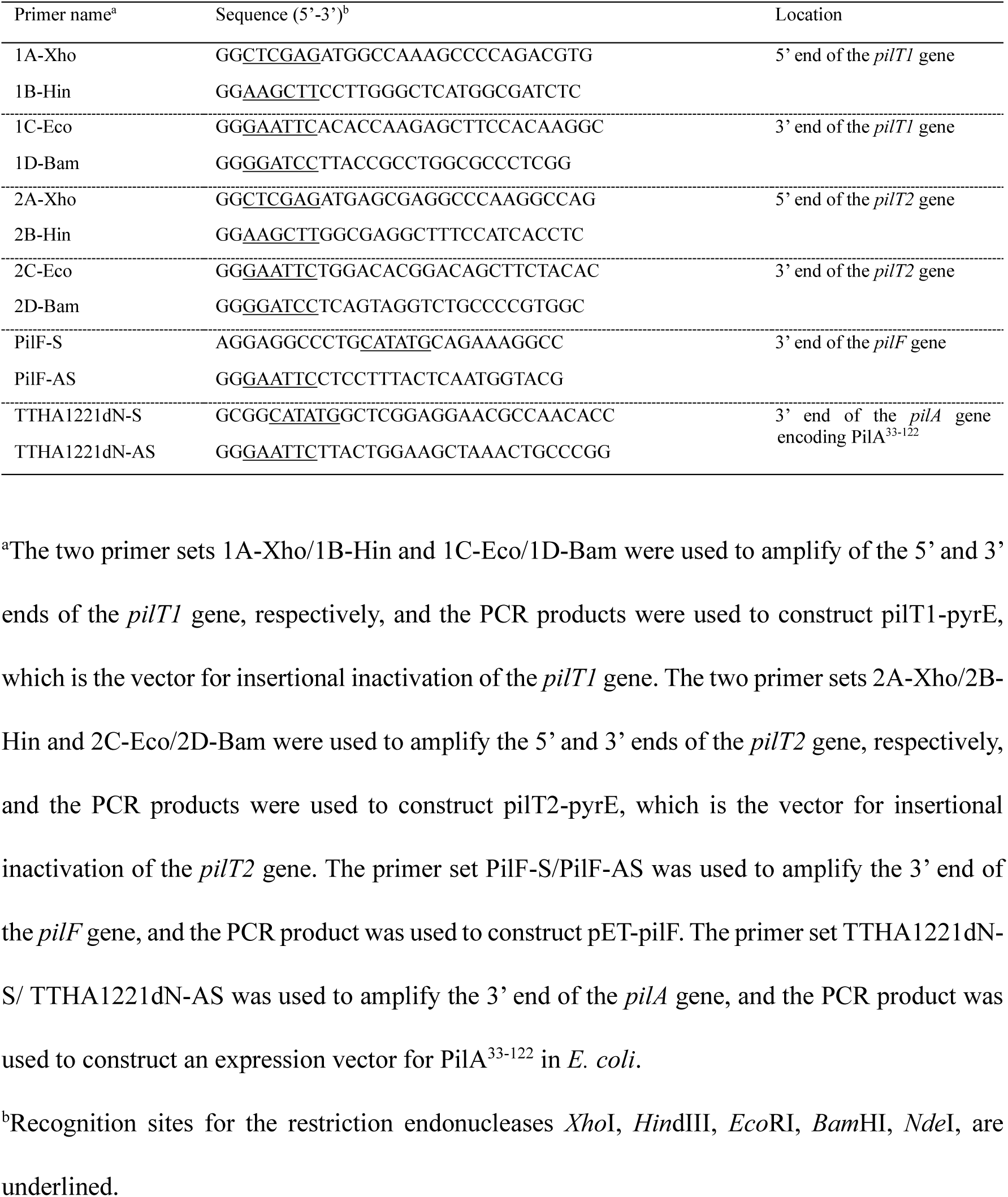
Oligonucleotides used in this study.

**Table S5.** Composition of Motility buffer.

| Strain | NaCl (mM) | pH |
| --- | --- | --- |
| *Thermus thermophilus* HB8 | 35 | 8.0 |
| *Thermus thermophilus* HB27 | 35 | 8.0 |
| *Thermus thermophilus* HB5002 | 35 | 8.0 |
| *Thermus thermophilus* HB5018 | 35 | 8.0 |
| *Thermus thermophilus* HC11 | 35 | 8.0 |
| *Thermus thermophilus* AK1 | 35 | 7.0 |
| *Thermus thermophilus* TMY | 35 | 7.5 |
| *Thermus aquaticus* YT-1 | 0 | 8.0 |
| *Thermus oshimai* SPS-17 | 35 | 8.0 |
| *Marinithermus hydrothermalis* T1 | 500 | 7.0 |
| *Meiothermus ruber* 21 | 0 | 7.5 |
| *Deinococcus soli* N5 | 0 | 7.0 |
| *Deinococcus grandis* KS0485 | 0 | 7.0 |
| *Deinococcus radiodurans* R1 | 0 | 7.0 |
| *Deinococcus rubellus* Ant6 | 0 | 7.0 |
| *Deinococcus radiophilus* RBD | 0 | 7.0 |

**Table S6.**
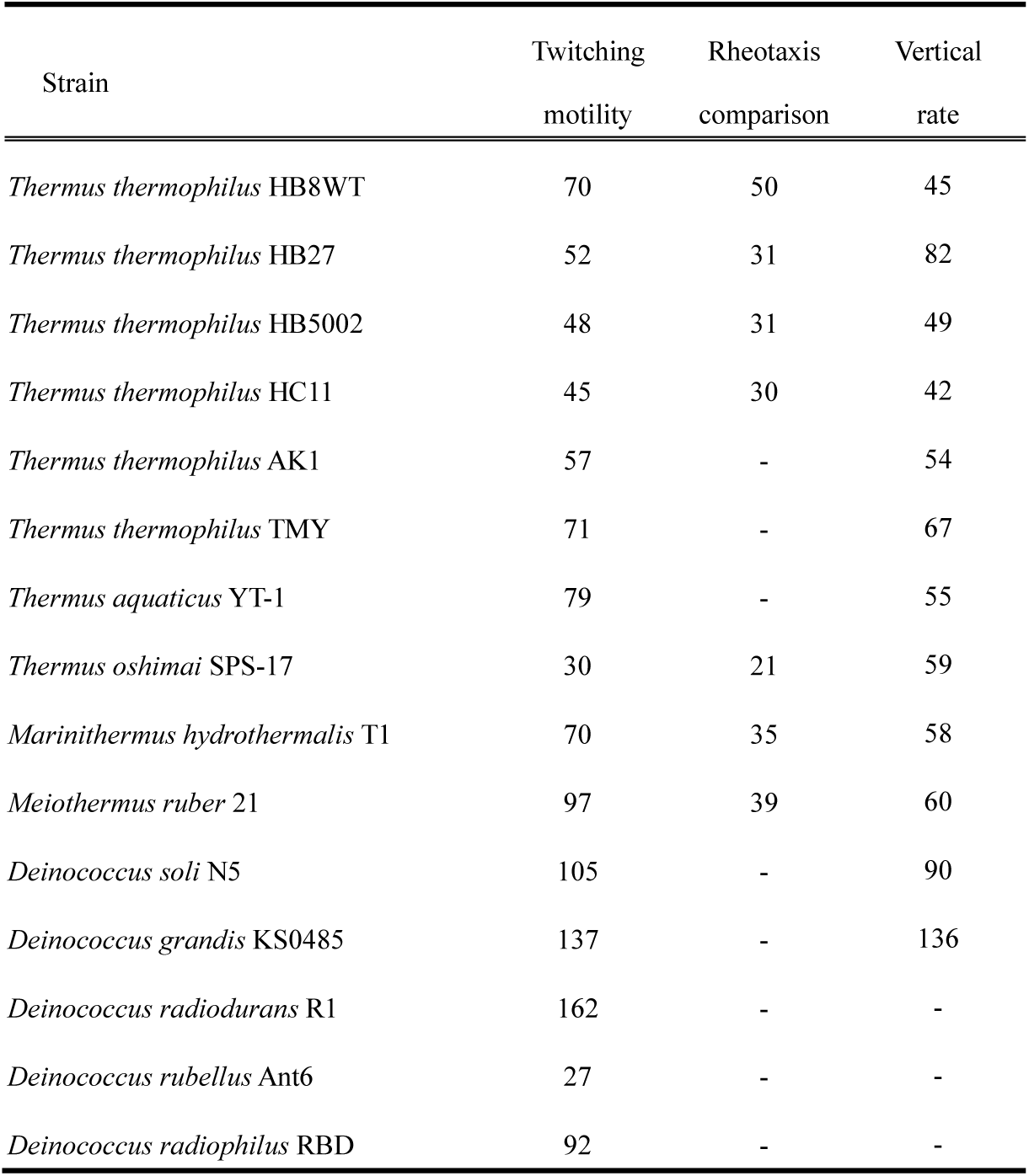
Sample size for the analysis of related species.

